# Complementary roles for hippocampus, anterior cingulate cortex, and orbitofrontal cortex in composing continuous choice

**DOI:** 10.1101/2025.03.17.643774

**Authors:** Assia Chericoni, Justin M. Fine, Taha S. Ismail, Gabriela Delgado, Melissa C. Franch, Elizabeth A. Mickiewicz, Ana G. Chavez, Eleonora Bartoli, Danika Paulo, Nicole R. Provenza, Andrew Watrous, Seng Bum Michael Yoo, Sameer A. Sheth, Benjamin Y. Hayden

## Abstract

Naturalistic, goal directed behavior often requires continuous actions directed at dynamically changing goals. In this context, the closest analogue to choice is a strategic reweighting of multiple goal-specific control policies in response to shifting environmental pressures. To understand the algorithmic and neural bases of choice in continuous contexts, we examined behavior and brain activity in humans performing a continuous prey-pursuit task. Using a newly developed control-theoretic decomposition of behavior, we find pursuit strategies are well described by a meta-controller dictating a mixture of lower-level controllers, each linked to specific pursuit goals. We find that hippocampal neurons encode the policy blending variable in a value-invariant manner and monitor policy switches after they occur. ACC neurons encode policy switches in a value-dependent manner, with value related modulation detectable several hundred ms before the switch, alongside a ramping increase in mean firing rate toward the switch. Meanwhile, OFC activity is consistent with an encoding of the current value structure of the task, rather than policy switching. Together these results are consistent with a tripartite functional division in which hippocampus serves as a controller over behavior, ACC serves as a meta-controller, and OFC provides a value context signal. Overall, our results shed light onto the complex processes associated with choice during naturalistic continuous interactive behavior.

## INTRODUCTION

Our understanding of how the brain implements reward-based choices is primarily based on studies of discrete choices with static options. Yet many of the decisions we face in the real world are continuous and interactive (Gordon et al., 2021; Maselli et al., 2023; Yoo et al., 2021). Consider a predator pursuing fleeing prey (Fabian et al., 2018; Fine et al., 2024; Yoo et al., 2020). At each moment, the predator can choose a single trajectory from a large continuous set of possibilities. That choice must *prospectively* take account of the fact that the prey will respond to its actions and *retrospectively* account for the instantaneous feedback it receives (Bertsekas, 1996; Stengel, 1994; Sutton & Barto, 2018). When there are multiple prey, the predator must also decide whether and when to adjust which goal, or blend of multiple goals, is pursued (Cisek, 2012; Gallivan et al., 2016; Gomi and Kawato, 1993; Haith et al., 2015; Sridhar et al., 2021;Yang et al., 2022). These changes in goal blend ratios are the closest analogue to choices in continuous contexts.

In continuous choice contexts, such as prey pursuit, there are multiple good options at the same time. A simple controller can guide the decision-maker to the best option under fixed conditions, but a more profitable policy may involve blending goals as the behavioral demands change. That would require a meta-controller to implement the optimal blending. In other words, choosing the best goal at any moment requires an action controller that can solve a non-trivial and nonlinear goal optimization problem (Bertsekas, 1996; Sutton and Barto, 1998). This poses computational challenges because a new nonlinear controller would have to be learned for each new set of goals experienced, and could not be generalized outside the goals it was designed around (Theodorou et al, 2010). A solution to this problem is the usage of compositional control, which builds on modern control theory, artificial intelligence and robotics (Dvijotham and Todorov, 2012; Peng et al., 2019; Haarnoja et al., 2024).

Optimal control theory has shown that the nonlinear control needed for solving continuous multi-goal decisions can be approximated via composition of individual linear controllers (Dvijotham and Todorov, 2012; Gomez et al., 2014; Pan et al., 2015; Peng et al., 2019; Wolpert and Kawato, 1998). In this framework, each individual controller is focused on resolving the best actions for a single goal, and the system’s behavior reflects a blended weighting of multiple controllers. A key benefit of *compositional control* is that a shared weighting mechanism can be generalized across environments with different goal-specific controllers (Gomez et al., 2014; Lake and Baroni, 2018; Matsuo et al., 2022; Todorov, 2009). A compositional controller, however, introduces a new problem, namely, the need for a meta-controller to adjudicate between controllers. The meta-controller should monitor both the environmental and internal shifts of goal priorities to determine the need for changes in the blending of individual controllers. Understanding if and how the brain might implement a flexible, compositional control strategy can elucidate the neural mechanisms enabling humans to arbitrate between competing goals in dynamic environments, and help us generalize choice to continuous contexts.

One implication of the blending hypothesis is that the brain should compute and explicitly represent the goal-specific policy weighting. Brain regions coding for a policy weighting should have preferential access to information needed for creating mappings of goals (e.g., goal distance) and goal specific control policies, and be able to compose them together (Kurth-Nelson et al., 2023; Whittington et al., 2022). This requirement closely aligns with a well-established role of the hippocampus in organizing knowledge into a cognitive map that can be referenced to guide flexible behavior (Tolman, 1948; O’Keefe & Nadel, 1978; Eichenbaum & Cohen, 2014; Behrens et al., 2018), and align with current task rule, context, or strategy (Kay et al., 2020; Behrens et al., 2018; Park et al., 2020; Sanders et al., 2019). Indeed, while hippocampal representations are strongly modulated by goals and task structure, value comparison and control-demand computations are more consistently attributed to frontal circuits (Rushworth et al., 2011; Wikenheiser & Schoenbaum, 2016; Shenhav et al., 2013; Behrens et al., 2018). This function is most akin to meta-control, which involves integrating information about task context, goal priorities, and expected outcomes to regulate control strategies (Yeung & Summerfield, 2012; Eppinger et al., 2021; Musslick et al., 2024). Previous work implicates the anterior cingulate cortex (ACC) in precisely these meta-control functions, including the monitoring and regulation of behavior across a wide range of decision-making contexts (Alexander & Brown, 2011; Shenhav et al., 2013; Kolling et al., 2016; Akam et al., 2021; Sarafyazd & Jazayeri, 2019; Shima & Tanji, 1998; Kennerley et al., 2006; Sheth et al., 2012). In particular, during decisions, ACC activity frequently exhibits ramping dynamics that precede choices and strategy switches, consistent with a role in accumulating evidence or integrating value-relevant signals to determine when a behavioral transition should occur (Blanchard et al., 2015; Hayden et al., 2011; Karlsson et al., 2012; Sarafyazd & Jazayeri, 2019; Kane et al., 2022; Balewski et al., 2023; Voloh et al., 2023). Building on this literature, we predicted that ACC neurons would encode policy switching in a value-dependent manner, showing anticipatory modulation several hundred milliseconds before an upcoming switch and a ramping increase in activity toward the time of the switch.

A meta-control system that reallocates control across goals also requires access to a representation of the task’s current value structure. The orbitofrontal cortex (OFC), which encodes the relative desirability of available options and the relationships between states and outcomes that define the value landscape of a task, can provide this information (Padoa-Schioppa & Assad, 2006; Schoenbaum et al., 2011; Hunt et al., 2018). Beyond scalar value signals, OFC has been proposed to organize this information into a cognitive map of task space, conceptually related to hippocampal cognitive maps but emphasizing value and outcome contingencies (Wilson et al., 2014; Wikenheiser & Schoenbaum, 2016). Consistent with this view, OFC representations may share structural similarities with hippocampal state representations, while differing in the information they emphasize. Thus, OFC provides the value representations over which meta-control mechanisms can operate. We predicted that OFC activity would primarily reflect the immediate value structure of the task, rather than the timing or dynamics of policy switching.

## RESULTS

Nineteen patients undergoing intracranial monitoring for epilepsy seizure onset zone localization, performed a joystick-controlled prey-pursuit task while we recorded brain activity from hippocampus, ACC, and OFC (**Figure 1A-D**, Chericoni et al., 2025). An additional cohort of fifty healthy participants also performed the same task, albeit without physiological recordings. On each trial, the participant used a joystick to move an avatar freely in a two-dimensional area to pursue fleeing prey (**Supplementary Video**). Avatar speed was determined by the magnitude of the joystick deflection from its center. On each trial, there were two prey that both evaded the participant. The prey were specified by different colors, indicating their value which changed along the dimensions of reward and speed (**Figure 1A-B**). Participants completed trials with three different combinations of prey value, ranging from no difference to a large difference in value (**Figure 1A**). A subset of participants (17 healthy participants and 3 patients with epilepsy) completed trials with a broader set of prey values spanning five levels (1-5). To ensure consistency across subjects, for these participants we remapped trials with a reward difference of 1 to 2, and trials with a reward difference of 3 to 4.

**Figure 1.**
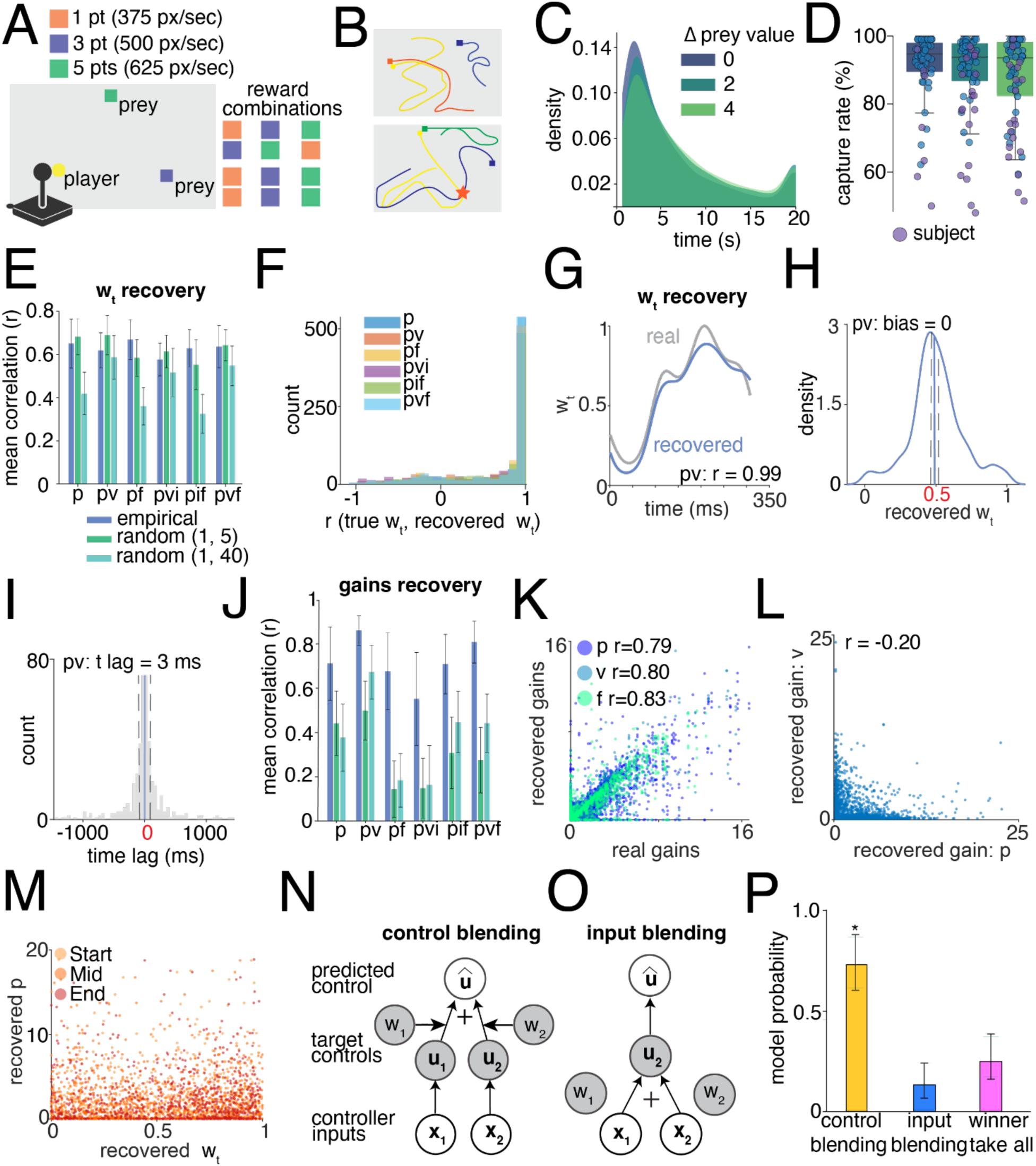
Continuous pursuit task and control choice model. (**A**) Schematic of pursuit task design. Prey were displayed as colored squares that denote their reward if caught and maximum speed. Participants’ avatar was displayed as a yellow circle, with avatar velocity determined by the magnitude of the joystick deflection from its center. On the right (reward combinations), each row represents a different trial type where the prey reward differed (three trial types). Participants also played trials in which prey of the same reward level were presented across all three unique prey types (three trial types). (**B**). The two schematics show example movement traces from two different trial types performed by a participant. (**C**). Kernel smoothing density (averaged across participants) of the time it took to capture a prey in trials with equal prey value (dark blue), with a small prey value difference (dark green) and large prey value difference (light green). The symbol Δ denotes the absolute difference between prey values. (**D**). Boxplot showing the capture rate median across trial types. Each dot represents one subject; blue dots denote healthy participants, purple dots participants with epilepsy (**E**). Barplot showing the recovery of w_t_ across simulations regimes and controller types, quantified as the Pearson correlation coefficient (r) between the simulated and recovered w_t_. Error bars indicate the standard deviation. (**F**). Histogram showing the distribution of Pearson’s correlation between the simulated and recovered w_t_ for the empirical simulation regime. (**G**). Example w_t_ trajectory from one simulated trial in the empirical regime for controller class *pv*. The grey trajectory indicates the empirically fitted w_t_ trajectory used as ground truth for simulation; while the blue trajectory shows the recovered w_t_ obtained by refitting the model to the simulated data. (**H**). Example histogram, for controller class pv in the empirical simulation regime, showing the distribution of the recovered w_t_ around the simulated w_t_, when w_t_ = 0.5 (switch point). The mean recovered w_t_ around the switch point was 0.5, resulting in zero bias, dashed lines represent 95% confidence intervals (CI), equal to [0.49, 0.52]. (**I**). Example time-lag histogram, for controller class pv in the empirical simulation regime. For each simulated trial, we computed the difference between the true time at which w_t_ crossed 0.5 and the time at which the recovered w_t_ crossed 0.5. The distribution peaks near zero, indicating minimal lag in recovered switch times. Dashed lines represent 95% CI, corresponding to [-80.5, 87.0] ms. (**J**). Barplot showing the recovery of controller gains across simulations regimes and controller types, quantified as the Pearson’s correlation r between the simulated and recovered gains. Error bars indicate the standard deviation. (**K**). Example scatter plot illustrating the correlation of the recovered controller gains vs simulated gains for the model including controller pvf in the empirical simulation regime, r denote Pearson’s correlation. (**L**) Example scatter plot illustrating lack of correlation among recovered gains for the model pv in the empirical simulation regime, r as in panel **H**. (**M**). Example scatter plot illustrating the lack of correlation between the recovered controller gains and the recovered w_t_ in the empirical simulation regime. w_t_ values are extracted at different time points within a simulated trial (start, middle, end). All Pearson’s r < 0.1. (**N**). Graphical model describing the control blending model. Shaded circles are latent parameters (w) or variables (u_1_,u_2_ = target specific controls). Transparent circles are measured variables, including inputs (x_1_,x_2_) and participant control signals (u^). (**O**). Graphical model describing the input blending model. (**P**). Posterior Bayesian model probability of models fit to subject data for control blending, input blending, and winner-take all models. The control blending was a significantly better fit across participants (*p* < 0.001 denoted by *).

Each trial ended with either the successful capture of one of the prey or after 20 seconds. Healthy participants contributed an average of 227 ± 8 (mean ± sd) number of trials each (Δprey value = 0: 78 ± 4 trials; Δprey value = 2: 101 ± 2 trials; Δprey value = 4: 47 ± 2 trials); while participants with epilepsy contributed an average of 81 ± 9 trials each (Δprey value = 0: 41 ± 5 trials; Δprey value = 2: 44 ± 4 trials; Δprey value = 4: 21 ± 2 trials). Thus, although the number of participants with intracranial monitoring is necessarily limited by the availability of human iEEG recordings, the analyses are based on tens of thousands of behavioral samples per subject.

Overall, we observed that as the difference in prey value increased, the trials also took longer (Δprey value = 0: mean t = 6.0 ± 0.08 seconds; Δprey value = 2: mean t = 6.9 ± 0.07 seconds; Δprey value = 4: mean t = 7.2 ± 0.1 seconds; all p < 0.05, t-test across value conditions) as anticipated, because the larger value prey moved faster, increasing the difficulty, **Figure 1C**. Across participants, mean capture rates were 90.8 ± 1.6% (mean ± sd) in the equal-reward condition, 89.0 ± 1.6% when prey values differed by the smallest amount (Δprey value = 2), and 87.6 ± 1.6% in the largest difference condition (Δprey value = 4; **Figure 1D**).

### Inferring continuous choice strategies

In discrete decision-making tasks, assessing a participant’s choice between options is trivial. In continuous decision contexts, inferring a participant’s moment-to-moment decision strategy is quite a bit more challenging. We need to estimate how their control signal (in this case, avatar velocity) is driven by one or more goals, in this case, by either a single prey or mixture of prey. One difficulty with identifying these mixing strategies is that participants may use a variety of low-level control strategies for pursuing individual targets. For example, participants may opt to minimize positional error, velocity error, integral of position error, estimated future position error, or any combination of these. The control strategy is unknown to the experimenter. Therefore, we must estimate both the participant’s per-target control strategy and the mixture of controllers.

The problems of estimating low-level controllers and target mixing are interrelated. To identify the target mixing at every point in time, we must estimate both the participant’s target specific controller and the mixture weight that combines these target controllers (w_t_; see **Methods** eq. 2-3). For the case of two possible pursuit targets, w_t_ encodes the continuum from fully pursuing one target (w_t_ = 0), to fully mixing targets (w_t_ = 0.5), to fully pursuing the other target (w_t_ = 1). w_t_ determines how much each target in our task contributes to the participant’s momentary control signal. If instead of estimating the controllers, however, we only used current position error controllers in tandem with estimating w_t_, this would provide incorrect estimates of w_t_ if they are using a different control strategy (e.g., current and future position error). To overcome this problem, we developed a model-based decomposition that leverages optimal control and compositionality theory to estimate these controllers and their mixtures (Todorov, 2009).

For each trial, we estimated w_t_ by modelling the participant’s joystick control using a second-order linear dynamic system (i.e., acceleration; see **Methods**). This process involved optimizing separate controller gains for each target simultaneous with optimizing w_t_. We repeated this process for all types and combinations of controller classes (see **Methods**), ranging from the usage of current position errors between cursor and target, to integral error, or one step ahead future position error (see **Methods** for a list of all classes). We used a Bayesian model averaging framework to recover the per trial estimate of w_t_. This involved weighting w_t_ for each controller class combination by the posterior probability of that fit.

### Validation and identifiability of the blended control model

To validate our modeling approach, we performed recovery of ground-truth simulated w_t_ signals and controller gains, and found that the model could accurately recover w_t_ across controller classes and complexity. Specifically, to ensure that recovery performance was evaluated both within and beyond the parameter range relevant to our data (Wilson & Collins, 2019), we repeated the recovery analysis across three gain regimes: random draws between 1-5, an extended range between 1-40 capturing a wide parameter space that encompasses the empirical regime (∼ 0.01-25), and an empirical range matching the parameters obtained by fitting the model to our dataset (including both empirical gains and empirical w_t_ trajectories). Recovery fidelity was quantified using Pearson correlation coefficients (r) between true and recovered parameters, averaged across runs (30 trials over 30 simulations). Across all simulations and controller classes, w_t_ recovery was strong on average (mean r = 0.57 ± 0.10, **Figure 1E**), and the distribution of such correlation was strongly skewed toward 1 (**Figure 1F-G**). Nonetheless, high correlations can hide systematic biases in recovery. Because our subsequent analyses build on the recovery of the switch points (w_t_ = 0.5), we compared the recovered w_t_ with the true w_t_ at switch points and measured their temporal lag. To prevent possible artifacts in the recovery estimates, we excluded trials in which the model output was saturated (w_t_ = 0 or 1; ∼9% of trials per simulation), although follow-up analyses showed that including them had a negligible effect on the results. When simulations were run using empirically fitted gains, the model was on average unbiased across controller classes (mean w_t_ = 0.49, 95% CI [0.47,0.51], **Figure S1A**).

Temporal lags between real and recovered w_t_ were minimal, with a median of 3.0 ms (95% CI [–96.4, 102.4] ms), well within one monitor display frame (that is, 16.67 ms, **Figure S1B**). Our preferred model pv, that is, the one that best fitted participant behavior, was unbiased (mean w_t_ = 0.5, 95% CI [0.49, 0.52], **Figure 1H**) with temporal lags close to zero (median lag = 3.0 ms, 95% CI [-80.5, 87.0] ms; **Figure 1I**). In simulations based on random or extended gain regimes, we observed a small negative bias (mean w_t_ = 0.48, 95% CI [0.45–0.50]; **Figure S1C, E**). Here too, median time lags remained below one frame (–11.8 ms, 95% CI [–134.4, 110.8] ms; **Figure S1D, F**), except for a few controller classes that showed shifts of up to two frames.

Correlation between real and recovered controllers gains was highest for the position-velocity (pv) controller class (pv: r = 0.67 ± 0.11; **Figure 1J** & **S2A**). For the remaining classes, the average recovery correlation was lower (r = 0.42 ± 0.15, **Figure1J-K** & **S2A**). Beyond recovery accuracy, we next assessed whether jointly estimating controller gains and w_t_ introduced identifiability issues. To examine whether the joint estimation of w_t_ and gain parameters introduced any identifiability issue (Wilson & Collins, 2019), as smaller w_t_ values could, in principle, compensate for larger gains and vice versa, we computed their correlations at the beginning, middle, and end of each simulated trial. These correlations were consistently low and stable across trial epochs (Start: r = 0.02 ± 0.16; Mid: r = -0.002 ± 0.14; End: r = 0.007 ± 0.14; **Figure 1M**), with no systematic differences across gain regimes (random or empirical) and controller classes (**Figure S3**). This pattern indicates that compensatory interactions between w_t_ and gain parameters were absent and both sets of parameters were robustly identifiable.

Because similar identifiability issues could also arise within mixture of controller classes, where different gain terms might compensate for one another, we repeated the analysis by correlating the recovered gains across error terms. For example, in the pv controller, we measured correlations between the recovered position and velocity gains. These correlations were generally modest and unsystematic across controller types, indicating that compensatory effects were minimal and that parameters were identifiable overall (r = 0.10 ± 0.04, **Figure 1L** and **Figure S4**). However, for the pvf model we observed higher correlations between velocity and force gains (mean r = 0.41 ± 0.20, **Figure S4**), and the velocity and position gains (mean r = 0.43 ± 0.20; **Figure S4**), suggesting that as model complexity increases, some degree of trade-off may occur. This effect does not impact our results, as in all participants the best-fitting model included only position and velocity errors (mean across simulation regimes r = 0.12 ± 0.29; **Figure 1L** and **S4**).

Lastly, we also found w_t_ is largely unrelated to subject speed (r = 0.14 ± 0.18, p = 0.38), relative distance (r = 0.11 ± 0.06, p = 0.53) or relative speed (r = 0.09 ± 0.02, p = 0.25) of the two prey across subjects. Importantly, these results indicate the controller mixing is identifiable and is a unique construct that is not just a reflection of a singular state variable such as relative distance.

We compared the fit of the blended controller model to participant behavior against two other potential strategies: (1) all-or-none target selection (i.e., winner-take-all) or (2) a blending strategy that mixes target input (e.g., relative position to subject) and uses a single controller. Notably, the all-or-none strategy is nested within the blending model, but the input blending model is distinct (**Figure 1N-O**, see proof in **S7**). Using simulations and ground-truth model recovery, we found a low confusion rate across models (correct classification for blending: 93.2%, all-or-none: 85.8%, and input mixing: 89.2%). We next evaluated fits of participant behavior with a Bayesian model comparison, with per trial and per subject model fits. The model posterior probabilities were used to adjudicate between the propensity of each strategy type. Model comparison indicated all subject’s behavior was best captured by a blended controller strategy (84% ± 8%, posterior model probability across subject; **Figure 1P**).

### Behavioral signatures of continuous pursuit strategies

We used the recovered w_t_ trajectories to understand participants’ pursuit behavior (example in **Figure 2A**). To visualize w_t_ behavior, we plot subjects’ averaged distributions across trial types, including trials with equal and different prey value (**Figure 2B**). If we define the pursuit of the more rewarding targets as moments when w_t_ > 0.85 and pursuit of the lower reward targets as moments when w_t_ < 0.15, we see the expected behavior. Participants spent a larger portion of time pursuing the more rewarding target in the small (t = -4.08, *p* < 0.0001) and large (t = -2.39, *p =* 0.02) reward difference trials. The remainder of the time (43%), pursuit behavior was spent in a blended state (0.15 < w_t_ < 0.85, **Figure 2C**). In trials with equal reward, as expected, subjects spent equal time chasing any of the prey (t = -0.93, p = 0.36) and 40% of the time in a blended state (**Figure 2C**). In summary, participants pursued the more rewarding target more often as the reward difference increased, otherwise they hedged their actions between controllers, acting in accordance with a softmax weighted decision strategy.

**Figure 2.**
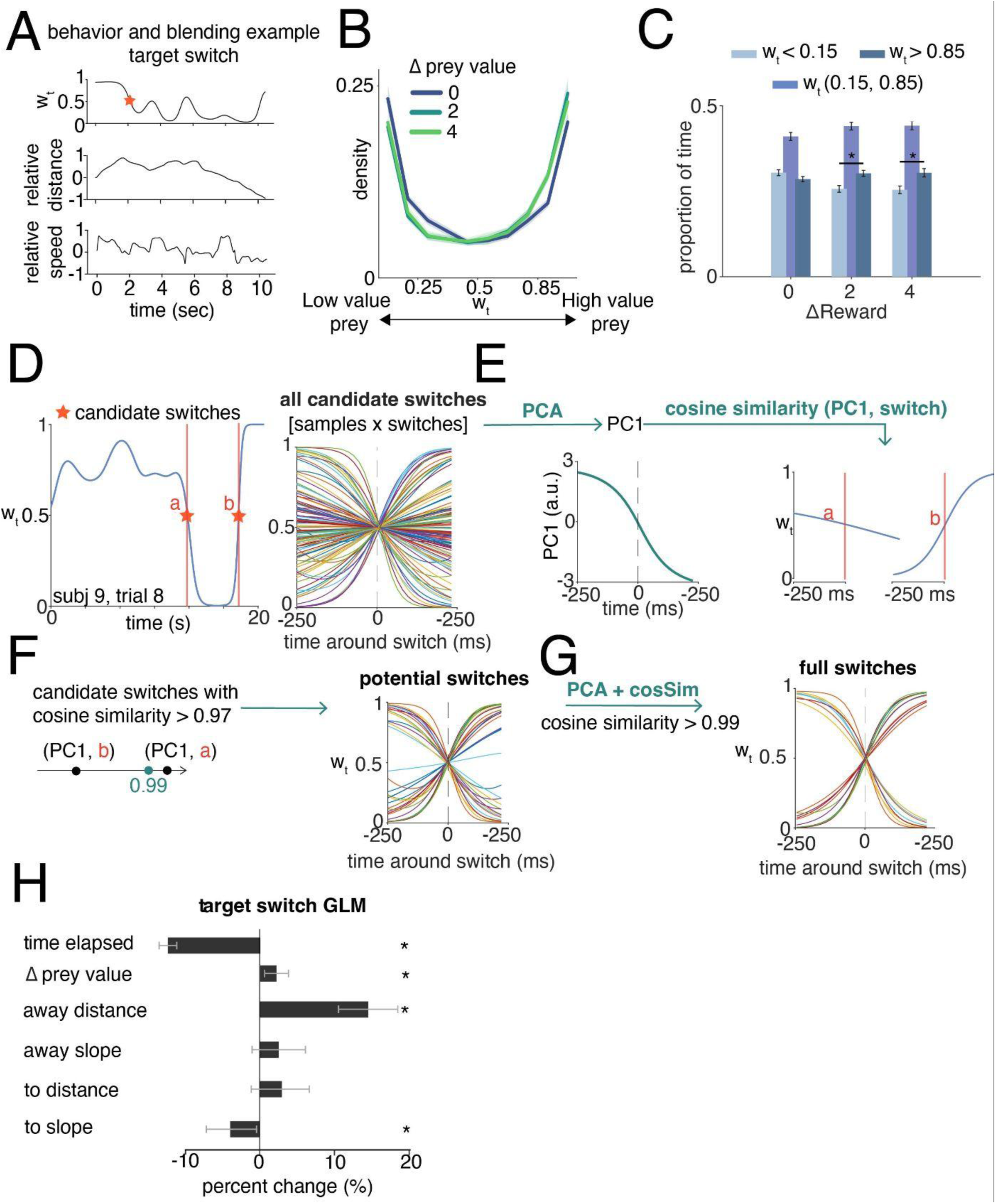
Target switch characterization. (**A**). Example time series from a single trial, showing the w_t_ trajectory (top), relative prey distance to the participant (middle), and relative prey-participant speed (bottom). The red star in the w_t_ trace marks an example candidate target switch. (**B**) Kernel smoothing density showing the distribution of controller blending (w_t_) for equal (blue), small (dark green) and large (light green) prey reward difference. The plot shows that as w_t_ is greater than 0.5, control is slightly biased toward the larger reward target. (**C**). Barplot showing the proportion of time spent pursuing each target or in a blended state, across reward conditions. (**D**) Schematic (real data) of all the candidate target switches as points where w_t_ crossed 0.5. (**E**) We used PCA on the candidate switches to find the 1st PC which captures the sigmoidal shape of the full switch trajectory. (**F**). We computed the cosine similarity between the 1st PC and all the candidate switches and retained those that crossed a threshold of 0.99. (**G**). We repeated the same process for a second time and retained only full switches. Sigmoids that go from high values to low values, represent switches from the lower value prey to the higher value prey (and vice versa) (**H**). Coefficients from hierarchical GLM predicting % change in odds of switching prey (significant predictors denoted by * with posterior probability > 0.95).

### Multiplexed states underlie goal switching

To determine which factors drove switches between targets, we first identified target switches (**Methods**), e.g., time series of switching from w_t_ < 0.15 to w_t_ > 0.85 (**Figure 2D-G**). Across subjects, this procedure yielded 13,357 candidate switches, of which 48% (6,366) were classified as full switches by our detection algorithm (**Methods**; **Figure 2D-G**). Of these full switches, 30.5% occurred in the equal-value condition, with the remainder distributed across unequal-value conditions (46.3% for Δprey value = 2; 23.2% for Δprey value = 4). Patients with epilepsy contributed 119 switches in the equal value condition and 798 in the different value conditions. The proportion of switches from the lower value prey to the higher value prey (48%) was comparable to the proportion of switches in the opposite direction (52%). Likewise, we found that, at switch point, the mean distance to the current (normalized distance for high > low switches: 0.76; while for low > high switches: 0.79; ttest, p = 0.30) and future targets (normalized distance for high > low switches: 0.79; while for low > high switches: 0.76; ttest, p = 0.29) did not differ across switch direction, indicating similar catchability.

We used a hierarchical Bayesian logistic model to determine factors that lead to changes in propensity for target switching (% change in odds of switching; **Figure 2H**). Increased distance from the pursued prey increased the likelihood of switching (14.6%), along with a small effect from differences in prey value (2.3%). The speed at which the switched-to target was moving away from the participant and the time spent pursuing the same target decreased the likelihood of switching (−4.1% and -12.4%; **Figure 2H**). Switching behavior was also driven by complex interactions among predictors, among them the interaction between the distance from the switched-to target and differences in prey value (13%).

These results show that target switches reflected a combination of multiple spatial, temporal and reward variables, rather than a single dominant factor.

### Controller blending is differentially encoded in ACC, hippocampal and OFC neurons

To examine the neural basis of continuous choice and w_t_, we recorded neurons in the ACC (n = 254 neurons; **Figure 3A**), hippocampus (hpc, n = 546 neurons, **Figure 3A**) and OFC (n = 103 neurons; **Figure 3A**). We characterized neural tuning to w_t_ by fitting neuron spikes with a Poisson generalized additive model (**Methods**) that uses a spline basis to estimate the tuning curves to w_t_, alongside auxiliary task parameters (e.g., relative prey value, relative prey distance, relative prey speed, avatar speed, trial time elapsed). As our main goal was estimating whether neurons exhibited disentangled (linear) tuning to w_t_ or multiplexed with prey value, we included both w_t_ and its interactions with prey value as predictors (**Figure 3B-C**). To ensure interpretability, we computed pairwise correlations and variance inflation factors among predictors (all < 2), confirming that collinearity was minimal and did not affect the analysis (**Figure 3D-E**). We found neurons were tuned linearly to w_t_ in all the brain regions (16.9% in ACC; 17.2% in hippocampus, and 39.8% in OFC, **Figure 3C**). We also found a substantial portion of neurons exhibiting nonlinear mixing of w_t_ with relative prey value (14.2% in ACC; 14.8% in hippocampus, and 35.0% in OFC, **Figure 3C**). Besides w_t_, we also found neurons tuned to all the auxiliary variables (**Figure 3C**).

**Figure 3.**
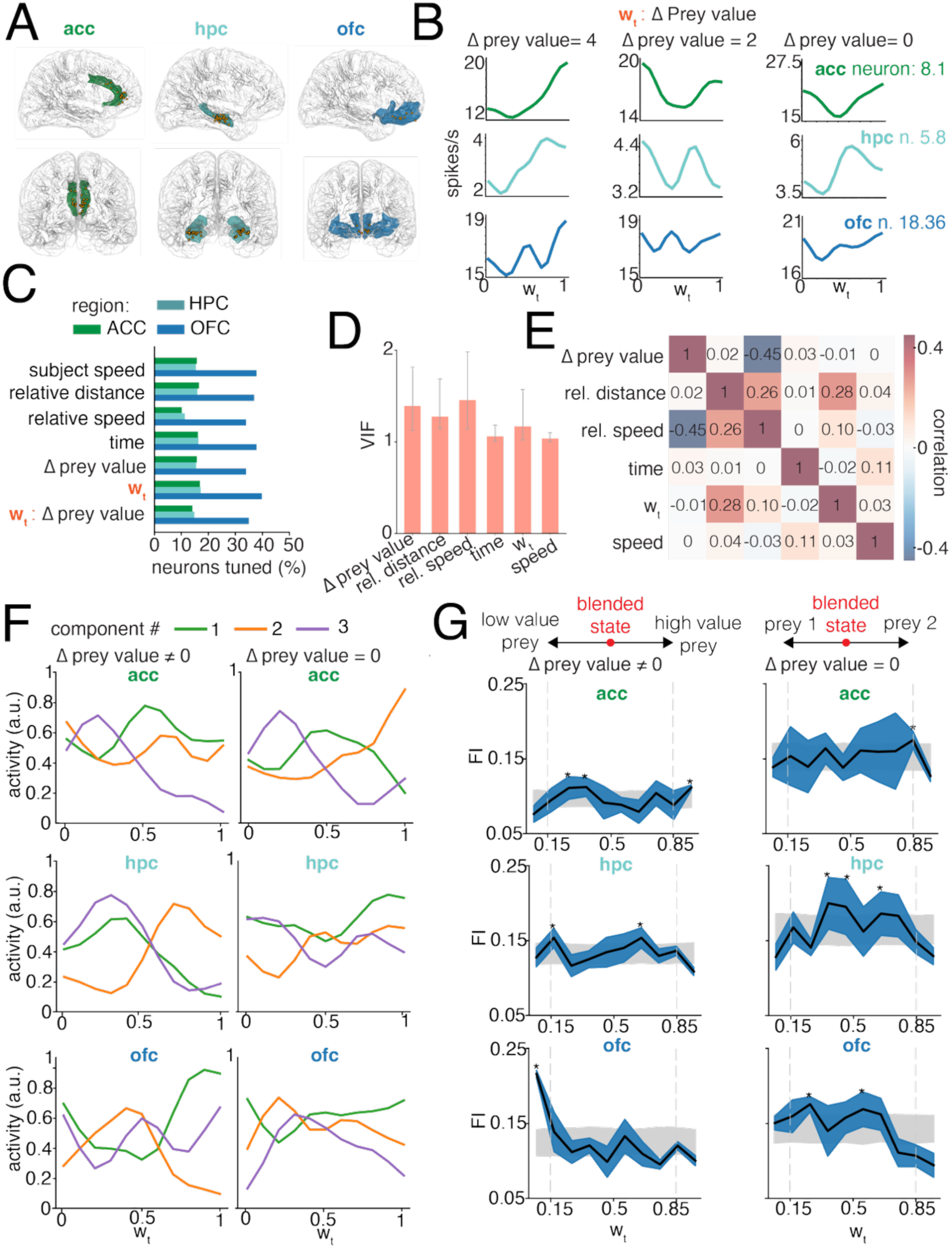
Neurons in the ACC, hippocampus and OFC are differentially tuned to w_t_. **(A)**. Recording sites from the ACC (left), hippocampus (center) and OFC (right). Dots represent electrodes across 19 patients. (**B**). Exemplar tuning curves to w_t_ across reward conditions (columns) and brain regions (rows), estimated with the Poisson generalized additive model (GAM). (**C**). Percent of neurons tuned to each variable across brain regions, estimated with the GAM. (**D**). Variance inflation factors (VIF) assess multicollinearity among predictors in the GAM. All VIFs are lower than 2, indicating minimal multicollinearity and stable coefficient estimates. Bars indicate mean VIF; error bars indicate the range across patients (**E**). Pairwise Pearson’s correlations among predictors, showing low to moderate correlations. (**F**). Non-negative matrix factorization components showing low-dimensional tuning bases to w_t_ for each brain region (rows) and reward condition (columns). (**G**). Cross-validated Fisher information with mean (black line) and standard error (blue area). Gray shading represents the permuted distribution with 95% confidence intervals. Significant information is shown per bin with an asterisk. Rows and columns are in panel F.

An important question is whether these brain regions differ in terms of their population tuning distribution to w_t_ across prey value conditions. To compare how the ACC, hippocampus and OFC represent w_t_, we used Fisher information (see **Methods**) because it quantifies how well a neural tuning distribution can discriminate small changes in w_t_. For example, if a population is chiefly concerned with the value of a pursuit strategy, the Fisher information will be skewed toward certain w_t_ values (e.g., pursuing higher-value prey). Conversely, a more similar Fisher information around low and high w_t_ indicates a more balanced representation of pursuing different prey. By comparing Fisher information across low and high w_t_ values, we can pinpoint whether each brain region’s tuning to w_t_ is more directed at coding for pursuit value (skewed Fisher information) versus faithfully coding for w_t_ (similar low and high Fisher information).

We first visualized the population tuning to w_t_ using a non-negative matrix factorization for dimensionality reduction (**Figure 3F**). Inspection of the ACC components points to a coding skewed towards high value prey (w_t_ > 0.85), especially in the different prey value condition (**Figure 3F**). Hippocampal components showed distributed tuning across w_t_, with similar population structure for low (w_t_ < 0.15) and high value prey (w_t_ > 0.85) across reward conditions (**Figure 3F**). Lastly, OFC components differentiated low and high w_t_ when prey values differed, with weaker differentiation in the equal value condition (**Figure 3F**).

We next quantified this by examining Fisher information in separate permutation tests for each of the low and high w_t_ levels. Confirming visual intuition (**Figure 3F**), for ACC we found significant w_t_ information for high prey values across reward conditions (permutation test, p < 0.005); while information in the blended state was significant only when prey values differed (**Figure 3G**). In the hippocampus, Fisher information did not preferentially emphasize extreme w_t_ values, but instead was high for blended states (0.15 < w_t_ < 0.85), particularly in the equal prey value condition (permutation test, all p < 0.005, **Figure 3G**). In OFC, Fisher information peaked at low w_t_ values when prey values differed (w_t_ < 0.15, permutation test, p < 0.001), while in the equal value condition, it was distributed across blended states (0.15 < w_t_ < 0.85, permutation test, p<0.001, **Figure 3G**).

These results suggest that the hippocampus implements a policy blending code that persists even when value asymmetries are removed, as reflected by similar representations for low and high value prey and increased blending representation across reward conditions. In contrast, ACC representations are skewed toward high value prey and selectively encode blended states only when prey values differ, consistent with a value dependent role in modulating the pursuit strategy. OFC shows sensitivity to distinct policy states in a value dependent manner, preferentially emphasizing the lower value (but more catchable) option when prey values differ.

### Neuronal dynamics track the switch dynamics

If a brain region encodes the controller w_t_ then neural population dynamics should exhibit a close parallel to behaviorally derived w_t_ dynamics (Khona and Fiete, 2022). In addition, if the neural coding of w_t_ is used to compose controllers, the neural population should convey a linear readout of w_t_ that is at least partially independent of different reward prey combinations.

To assess whether neural dynamics encode the w_t_ dynamics, we aligned neural activity to target switches because the w_t_ trajectory during target switches follows a canonical sigmoidal-like pattern (**Figure 4A**). This sigmoidal w_t_ provides a template to assess whether neural population dynamics track w_t_. To quantify the similarity in neural and w_t_ dynamics, we decomposed the neural populations aligned to target switches using demixed principal components analysis (dPCA; Kobak et al., 2016, **Methods**) and did so separately for each brain region (**Figure 4B**). The decomposition was designed to separate dynamics from two types of switches, switches from the low to the high-value target and vice versa (here we included only trials with different prey value, **Methods**). We visualized the dynamics by projecting the population activity for each area onto the top dPCA component related to the controller w_t_ and projected held-out test trials on the top component to quantify linearly decodable information about w_t_ (see **Methods**). With this analysis, we can ask if a brain region is coding for switch direction (from high to low value prey and vice versa).

**Figure 4.**
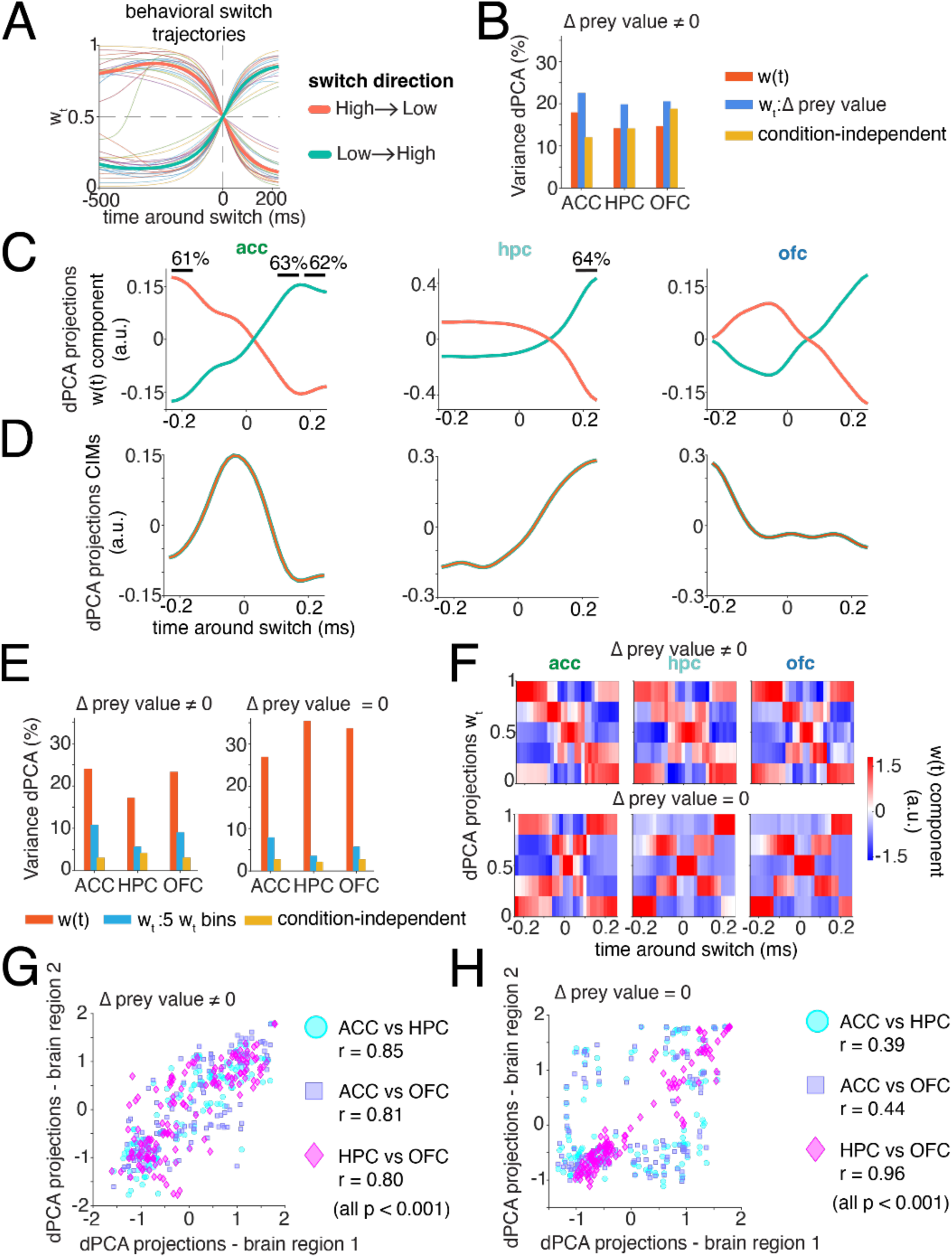
Low-dimensional controller blending dynamics during goal switch. (**A**). Controller blending dynamics extracted during target switches and aligned to switch time (**Methods** and **Figure 2D-G**). Orange traces denote switches from higher to lower value prey, and green traces denote switches from lower to higher value prey, averaged across participants. Shaded lines represent the averaged trajectories per participant. (**B**). Percentage of variance explained by each task factor from dPCA for each brain region. Here, dPCA was configured to separate switch types. (**C**). Neural population activity projected onto the leading w_t_ dPCA component for each brain region. Black dashed lines indicate time bins with decoding accuracy significantly above chance; values above the dashed lines report decoding accuracy. Green and orange traces are as in panel A. (**D**). Neural dynamics projections for condition independent modes (CIM). Green and orange traces overlap. (**E**). Percentage of variance explained by each task factor from dPCA for each brain region, with dPCA configured to separate w_t_ representations across five bins. Results are shown separately for the different prey value condition (left) and the equal prey value condition (right). (**F**). Neural population activity projected onto the leading w_t_ dPCA component for each brain region (columns) and reward condition (rows). Heat maps show mean dPCA projections across five w_t_ bins as a function of time relative to the switch (color indicates component magnitude, a.u.). (**G**) Pairwise correlations between dPCA projections across brain regions in the different prey value condition. Each point represents a matched time w_t_ bin across regions; values report Pearson correlation coefficients. (**H**). Same as panel **G**, but for the equal prey value condition.

We found that for ACC the top dPCA component for w_t_ explained 17.8% of the variance, while its interaction with prey value difference explained 22.5% of the variance (**Figure 4B**). ACC had significant linearly decodable switch direction at switch onset (61% decoding accuracy, permutation test: p = 0.03, **Figure 4C**) and at the end of the switch (62.5% decoding accuracy, p = 0.03, **Figure 4C**). Furthermore, the ACC neural dynamics closely mirrored the sigmoidal w_t_ trajectory predicted by behavior (Pearson r = 0.99, p < 0.0001). These results show that ACC activity is value dependent and potentially drives and monitors the switch decision.

We also found that, in the hippocampus, neural population dynamics are explained by the w_t_ dPCA component (14.2%), and its interaction with differences in prey value (19.9%, **Figure 4B**). Hippocampal w_t_ dPCA projections resembled the switch trajectories (r = 0.84, p < 0.001; **Figure 4C**), but this correlation was significantly weaker than in ACC (Fisher test for correlation, p < 0.001). Consistent with this, we could decode switch direction from hippocampal neurons only at switch offset (64% decoding accuracy, p < 0.001, **Figure 4C**). These findings suggest that the hippocampus reflects how controllers are mixed and updates the policy representation at switch offset.

Similarly, in OFC, neural population variance was primarily explained by the w_t_ dPCA component (14.7%) and its interaction with differences in prey value (20.6%, **Figure 4B**). The correlation between the w_t_ dPCA projections and switch trajectories was comparable to that observed in the hippocampus (r = 0.86, p < 0.001, **Figure 4C**) and significantly lower than in ACC. Despite this correspondence, switch direction could not be decoded from OFC activity at any time point. Notably, OFC projections converged at switch onset and diverged only after the transition, suggesting that OFC neurons have a value representation of the selected policy only at the end of the switch.

The dPCA also indicated substantial condition-independent modes (CIM) in all brain areas (**Figure 4D**). The CIMs represent w_t_ evolution over time independently of value. The top CIMs explained 12.0 % for ACC variance, 14.1% for hippocampus, and 18.8% for OFC (**Figure 4D**). While the origins of these CIM dynamics are unclear, their profiles indicate distinct temporal patterns associated with the progression of switch decisions across the three brain regions.

Next, we examined how neural representations of switch dynamics differ between the equal and different prey value conditions. To do so, we re-applied dPCA, this time configuring the decomposition to isolate w_t_ dynamics across five discrete w_t_ levels (five w_t_ bins, **Figure 4E**). This decomposition was applied separately to trials with different prey value and trials with equal prey values (**Figure 4E**) and performed independently for each brain region. We found that across brain regions and reward conditions, most of the variance was explained by the time-dependent w_t_ components (variance explained 1st w_t_ PC. Δprey value ≠ 0, ACC: 24.0%; hippocampus: 17.2%; OFC: 23.3%; Δprey value = 0, ACC: 26.9%; hippocampus: 35.4%; OFC: 33.7%). Projecting population activity onto the top dPCA component related to w_t_ revealed differences in how policy dynamics were structured by value differences (**Figure 4F**). When prey values differed, ACC w_t_ projections exhibited an asymmetric structure across the five bins, forming two distinct diagonals in the heatmap, which collapsed into a more symmetric pattern when prey values were equal (**Figure 4F**). This confirms that ACC policy representations are shaped by value asymmetry. In contrast, hippocampal activity symmetrically spanned the full w_t_ continuum in the different-value condition but shifted toward blended control states when values were equal, consistent with a value-invariant representation of the policy blending code (**Figure 4F**). OFC projections were largely symmetric across reward conditions and continued to emphasize w_t_ extremes even when prey values were equal (**Figure 4F**). Correspondingly, correlations between both hippocampal and OFC projections with ACC were reduced in the equal-value condition (Fisher test, p < 0.001; **Figure 4G-H**), whereas correlations between hippocampus and OFC increased.

### Neural ramping during goal switching

Previous studies examining discrete decisions indicate ACC population activity often exhibits ramping dynamics aligned to the onset of decisions (Balewski et al., 2023; Sarafyazd and Jazayeri, 2019). Building on these findings, we hypothesized that ACC may also play a role in driving target switch decisions exemplified by ramping signals coinciding with the target switch onset (**Figure 5A**), while no such ramping should appear during time windows without switches. Furthermore, previous studies of the neural decision-making processes often find ramping profiles at the single neuron level (e.g., Gold and Shadlen, 2007; Kiani et al., 2014; Luo et al., 2024). Therefore, we asked whether switch aligned ramping dynamics observed at the population level might also be evident within subpopulations of single neurons.

**Figure 5.**
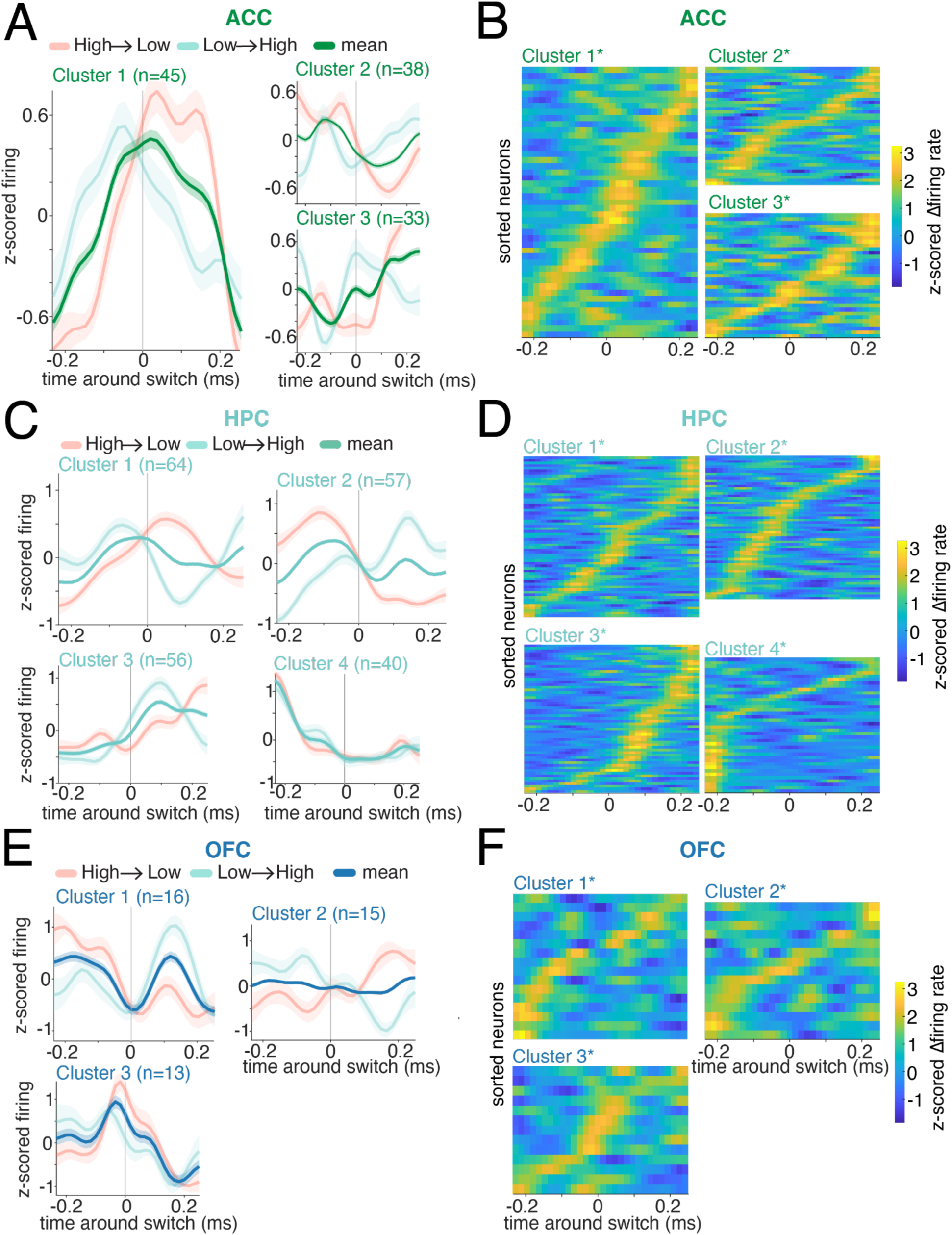
Neural ramping during changes of mind. **(A).** Mean cluster firing rates in ACC time locked to switch onset. The green trace represents the average activity across switch directions (high > low; low > high) (**B)**. ACC z-scored cluster selectivity, neurons are sorted by the switch-aligned time of maximal difference between switch and control activity. (**C**). Mean cluster firing rates in hippocampus, time locked to switch onset. The light green trace represents the average activity across switch directions. (**D**). Hippocampal z-scored cluster selectivity, as in panel (**B**). (**E**). Mean cluster firing rates in OFC, time locked to switch onset. The dark blue trace represents the average activity across switch directions. (**F**). OFC z-scored cluster selectivity, as in panel (**B**).

We used a method that combines k-means clustering with linear dynamic systems, which allows us to cluster neurons based on their dynamics (**Methods**). The number of clusters was optimized (silhouette score) separately for each brain region, and only clusters exhibiting significant selectivity for whether a target switch occurred were retained using a permutation test. This process yielded three clusters for the ACC (all permutation tests, p < 0.01, **Figure 5A-B**). Visually examining their temporal properties, we found that the largest cluster (cluster 1, n = 45) exhibited ramping activity that peaked at switch time and spanned the entire switch window (dark green trace, **Figure 5A**). Ramping associated with switches from higher- to lower-value prey was stronger and peaked later (33 ms). Notably, for both switch types this activity preceded the time at which switches became detectable in behavior (−150 ms; **Figure 4A**), indicating that ACC engagement emerges before the behavioral switch. Moreover, the sustained activity across the switch window is consistent with a monitoring role for ACC.

In the hippocampus we found four clusters of neurons (all permutation tests, p < 0.01; **Figure 5C-D**). Visually examining their temporal properties, we found that cluster one and three (and cluster 4 with inverted polarity) recapitulate the dynamics observed in population-level decoding (**Figure 4C**). Specifically, neurons in these clusters exhibited selectivity for target switches after switch onset rather than ramping toward the decision point (time = 0). Neurons in cluster 2, instead closely tracked the controller w_t_ over time, consistent with a representation of policy state rather than switch-related ramping (**Figure 5C**).

In OFC, we identified three significant neuronal clusters (all permutation tests, p < 0.01; **Figure 5E–F**), each exhibiting distinct temporal dynamics, without clear ramping toward the switch.

## DISCUSSION

Choice in continuous contexts poses multiple interrelated control problems (Merel et al., 2019; Yoo et al., 2021; Burge and Bonnen, 2025). We must combine dynamic feedback and predictions of future target states to decide which goals to pursue and how to pursue them. Here, we use control theory to understand continuous choice in a multi-prey pursuit task. We find that control is parsimoniously explained as being composed of a blend of discrete goal-specific control policies, and is implemented by at least three brain regions with complementary roles. Compositional control allows for efficient generalization across novel environments, such as different reward combinations (Driscoll et al., 2024; Johnston et al., 2024; Sandbrink et al, 2024). For example, when novel combinations of familiar prey are encountered during pursuit, there is no need to learn a new set of controllers, as existing ones can be readily combined.

Our modelling results in, essentially, an ethogram of behavior (Anderson & Perona, 2014). Previous studies have mostly used unsupervised and clustering methods to identify behavioral motifs (equivalent to controllers here) in naturalistic behavior (Calhoun et al., 2019; Voloh et al., 2023; Weinreb et al., 2024; Wiltschko et al., 2015). One benefit of the control theoretic approach, however, is the ability to detect blended controllers. The conventional approach, then, risks misinterpreting neural data that is derived from blended policies. Consequently, our results suggest that understanding ethological, high-dimensional behavior must go beyond simply categorizing discrete behavioral motifs.

The hippocampus has not historically been associated with control as much as it has been associated with mapping and with memory. Nonetheless, small but growing set of studies has extended its role to potentially include control functions, which may be integrated with its other roles (e.g., Stachenfeld et al., 2017; Vikbladh et al., 2019; Bakkour et al., 2019; Whittington et al., 2020; Edelson and Hare, 2023; Jensen et al., 2024). Although classically associated with spatial mapping (O’Keefe & Nadel, 1978), the hippocampus is now understood to represent relational structure, latent behavioral states, and structured cognitive maps that support flexible behavior across domains (Eichenbaum & Cohen, 2014; Gershman & Niv, 2010; Behrens et al., 2018). In navigation and pursuit tasks, hippocampal populations encode multiple agents, trajectories, and goals simultaneously, consistent with a role in maintaining parallel, control-relevant state representations (Aronov et al., 2017; Chericoni et al., 2024; Nieh et al., 2021), and these representations interact with prefrontal task representations to support flexible inference and planning (Wikenheiser & Schoenbaum, 2016; Behrens et al., 2018). This literature in turn suggested our core hypotheses in this study, namely, that hippocampus would play a role as a controller of behavior.

More broadly, we found neural results that align with and extend the theory that the hippocampus implements a cognitive map (Behrens et al., 2018; O’Keefe and Nadel, 1979; Zuthsi et al., 2025). The cognitive map theory posits that the hippocampus creates structured representations of the world and does so by composing and binding individual basis elements (Kurth-Nelson et al., 2023; Whittington et al., 2022). Previous studies have focused on hippocampal compositions over visual scene construction (Schwartenbeck et al., 2023), spatial information (Hoydal et al., 2019), or novel concept compounds (Barron et al., 2013). We extend these findings by showing the hippocampus also encodes compositions that bind goal-specific control policies. Specifically, hippocampal activity encodes the policy blending variable in a value-agnostic manner and primarily reflects the current policy state, with switch value information emerging only after the switch has occurred.

The present results indicate that ACC carries a robust meta-control signal, one that is directly associated with, and tracks, the process of switching from one controller to another. That control signal precedes the switch and appears to begin before the switch is detectable in behavior; it also shows a continuous ramping (in a latent space) in the leadup to the switch. This potential control-related role relates to a long tradition that emphasizes the role of ACC in control (Botvinick and Cohen, 2000; Schall, 2003; Shenhav et al., 2013; Heilbronner and Hayden, 2011; Paus, 2001). It is also consistent with ideas that extend this framework to meta-control (Verguts, 2017; Silvetti et al., 2018). These ideas in turn align with longstanding, albeit controversial, ideas that ACC serves as a conflict monitor, in that activity is highest at the time the need for control is maximized (Botvinick et al., 2001; Botvinick et al., 2004; Braver et al., 2001; Kerns et al., 2004; van Veen et al., 2001). The ramping effects, in turn, are reminiscent of a large literature demonstrating that ACC accumulates evidence for strategic shifts in decision-making or rule application (Hayden et al., 2011; Johnston et al., 2007; Kennerly et al., 2006; Kolling et al., 2012; Sarafyazd and Jazayeri, 2019; Shenhav et al., 2016). We also find that this ramping switch signal is dependent on the value of switching goals, a finding in line with previous studies and theories linking ACC to integrating cost-benefit information into decisions (Amiez et al., 2007; Friedman et al., 2015; Shenhav et al., 2016). A possible explanation of our result is that abandoning a high value versus low value prey represents decisions with different values and costs, thus eliciting different ramping signals. The implication is ACC is not merely signaling movement transitions but is integrating decision variables. This integration likely involves weighing the cost and benefits of switching targets, aligning with foraging models (Stephens & Krebs, 1986; Hayden et al., 2011; Kane et al., 2022).

Note that meta-control can be either implicit or explicit. Explicit meta-control would involve a part of the system that serves to monitor the success of the current meta-control strategy and modify it when needed. Implicit meta-control could involve, for example, competition between different controllers, such as mutual inhibition systems. These two arrangements can be difficult to disentangle in practice. One way to distinguish them is through neural data. Our data, which show clear explicit monitoring and meta-control signals in ACC, therefore provide evidence for the explicit implementation.

Our results suggest that OFC encodes the immediate value structure of the task, and has a minimal role in policy switching. A meta-control system that reallocates control across goals requires access to a representation of the task’s current value structure (Sutton & Barto, 1998; Daw et al., 2005; Botvinick et al., 2009). The orbitofrontal cortex (OFC) encodes the relative desirability of available options and the relationships between states and outcomes that define the value landscape of a task, core components of task structure (Padoa-Schioppa & Assad, 2006; Schoenbaum et al., 2011; Hunt et al., 2018). Beyond scalar value signals, OFC has been proposed to organize this information into a cognitive map of task space, conceptually related to hippocampal cognitive maps but emphasizing value and outcome contingencies (Wilson et al., 2014; Wikenheiser & Schoenbaum, 2016). Consistent with this view, OFC representations may share structural similarities with hippocampal state representations, while differing in the information they emphasize (Wilson et al., 2014; Wikenheiser & Schoenbaum, 2016). In this context, our data suggest that OFC provides the value representations over which meta-control mechanisms can operate.

We find a stronger ramp for the High > Low switch. This difference may reflect the additional demand for control needed when abandoning a larger potential reward and switching to a lower value one (cf. Hayden et al., 2011; Kennerley et al., 2006; Kolling et al., 2012; Shenhav et al., 2013; Heilbronner & Hayden, 2016). Furthermore, switching from the higher value prey represents an unexpected deviation from the default policy, which could enhance ACC engagement (that is, a stronger ramp). According to many theories, ACC activity is preferentially engaged during deviations from default behavior (Hayden and Platt, 2011; Kolling et al., 2012; Holroyd & Yeung, 2012). For example, the PRO model predicts increased ACC activity when an expected outcome fails to occur (Alexander & Brown, 2011; Brown & Alexander, 2017), consistent with studies linking ACC to deviations from default policies, control override, and evaluation of costly strategy switches (Botvinick, 2007; Kolling et al., 2012; Shenhav et al., 2013; Heilbronner & Hayden, 2016). Because catch likelihood is comparable for both prey in our data - proportions of high to low (52%) and low to high (48%) switches across subjects are comparable - the continuation of the expected behavior is mainly driven by value rather than ease of capture. Thus, switching away from the higher value prey represents the occurrence of the unexpected outcome, which may contribute to the stronger ramp for switches from high to low (Botvinick, 2007). In any case, these results clearly implicate ACC in the meta-control aspect in a way that hippocampus and OFC are not as clearly involved.

Our findings characterize continuous decisions as arising from the dynamic reallocation of control across competing goals. Rather than treating actions as independent choices, this perspective emphasizes their temporal dependence on ongoing control states. Our control-theoretic analysis of behavior identifies likely computational elements of continuous decision-making, while physiological analyses point to their potential neural substrates. In particular, our results indicate that three regions that play pivotal, and arguably distinct, roles in discrete choice also assume different roles when choice unfolds continuously over time, in ways that align with their known functions. Our results do not, however, imply that other regions play no role in continuous choice. For example, parietal cortex, which is closely associated with control of behavior in the peripersonal space (Milner and Goodale, 1995), almost certainly plays a major role, and dorsolateral prefrontal cortex likely plays an important role in regulating sensorimotor transformations (Miller and Cohen, 2001). We anticipate that future studies will delineate the full circuitry for continuous decisions.

## Acknowledgements

We thank Paul Schrater, Xaq Pitkow, Jon Pearson, and Scott Linderman for invaluable discussions and insights on solving the controller problems. We thank Jeff Johnston for discussion on neural analysis. We also thank Joshua Adkinson, Raissa Mathura, and Victoria Pirtle for invaluable assistance. This project was supported by NIH grants R01 DA038615, R01 MH125377, U01 NS121472, and R01 MH129439, and by the McNair foundation.

## Data availability statement

All data will be made available upon publication

## Code availability statement

All code is available at https://github.com/BCM-Neurosurgery/PreyPursuit

## METHODS

All procedures were approved by the Baylor College of Medicine Institutional review board.

### Human intracranial neurophysiology

Experimental data were recorded from 19 adult patients (12 males and 7 females) undergoing intracranial monitoring for epilepsy. The hippocampus, ACC, and OFC were not a seizure focus area of any patients included in the study. Single neuron data were recorded from stereotactic (sEEG) probes, specifically AdTech Medical probes in a Behnke-Fried configuration. Each patient had an average of 3 probes terminating in left and right hippocampus. Electrode locations are verified by co-registered pre-operative MRI and post-operative CT scans. Each probe includes 8 microwires, each with 8 contacts, specifically designed for recording single-neuron activity. Single neuron data were recorded using a 512-channel Blackrock Microsystems Neuroport system sampled at 30 kHz. To identify single neuron action potentials, the raw traces were spike-sorted using the WaveClus sorting algorithm (Chaure et al., 2018) and then manually evaluated. Noise was removed and each signal was classified as multi or single unit using several criteria: consistent spike waveforms, waveform shape (slope, amplitude, trough-to-peak), and exponentially decaying ISI histogram with no ISI shorter than the refractory period (1 ms). We only used single neuron activity for all analyses. Participants included in our study received a standard monetary compensation, which did not depend on task performance. All the participants received the same compensation.

### Healthy participants

In addition to the dataset with intracranial recordings, additional behavioral data were collected from 50 healthy adult participants who performed the same joystick-controlled prey-pursuit task but without physiological recordings. Of these participants, 33 were recruited at Baylor College of Medicine, and the remaining 17 were recruited at the University of Minnesota. All healthy participants provided informed consent in accordance with protocols approved by the respective institutional review boards at each site. Also here, participants received standard monetary compensation that did not depend on task performance.

### Experimental apparatus

To control their computer avatar, subjects used a joystick that was a modified version of a commercially available joystick with a built-in potentiometer (Logitech Extreme Pro 3D). The joystick position was read out in the MATLAB psychophysics toolbox running on the stimulus control computer.

### Task description

At trial start, two or three shapes appeared on a gray computer monitor placed directly in front of the subject. The yellow circle (15-pixels in diameter) was an avatar that represented the subject and began at the center of the screen. Subject position was determined by the joystick and was limited by the screen boundaries. A square (30 pixels in length) represented the prey(s). Prey movement was determined by a simple algorithm (see below). Each trial ended with either the successful capture of the prey or after 20 seconds, whichever came first. Successful capture was defined as any spatial tangency between the avatar circle and the prey square. Capture resulted in an immediate numeric reward.

Prey movement was generated interactively using an A-star algorithm. Specifically, for every frame (16.67 ms), we computed the cost of 15 possible future positions the prey could move to in the next time-step. These 15 positions were equally spaced on the circumference of a circle centered on the current position of the prey, with a radius equal to the maximum distance the prey could travel within one time-step. The cost was based on two factors: the position in the field and the position of the subject’s avatar. The field that the prey moved in had a built-in bias for cost, which made the prey more likely to move toward the center. The cost due to distance from the avatar of the subject was transformed using a sigmoidal function: the cost became zero beyond a certain distance so that the prey did not move, and it became greater as distance from the avatar of the subject decreased. From these 15 positional costs, the position with the lowest cost was selected for the next movement. If the next movement was beyond the screen range, then the position with the next lowest cost was selected until it was within the screen range.

The maximum subject speed was 23 pixels per frame (each frame = 16.67 ms). The maximum and minimum speeds of the prey varied across subjects and were set by the experimenter to obtain a large number of trials. Specifically, speeds were selected so that subjects could capture prey on no more than 85% of trials. To ensure sufficient time of pursuit, the minimum distance between the initial position of each subject avatar and prey was 400 pixels.

### Behavioral modeling

#### Overview of control choice decomposition

The problem we are studying here is how subject’s decide between how to pursue goals on a moment to moment basis. This means we must estimate a decision variable over how targets are selected to drive control actions. We motivate this problem by first considering the low-level, individual goal problem. For a given target, subject’s must deploy some policy that dictates their actions for reaching that goal, 𝑢_*k*_(𝑡). Here, we consider the controls as the output of a linear controller that is driven by a set of gains that dictate controls (library of controller types considered in Table 1). For example, the simplest proportional controller with a gain (𝑔_*k*_) that minimizes distance to target (𝑒_*k*_) for a single target 𝑘 is:

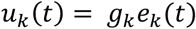

**Table 1.**

| Equation | Description |
| --- | --- |
| $u_{pred} = g_p e_p$ | Current position error |
| $u_{pred} = g_p e_p + g_v e_v$ | Current position and velocity errors |
| $u_{pred} = g_p e_p + g_f e_f$ | Current and future position errors |
| $u_{pred} = g_p e_p + g_i \int e_p e_p$ | Current and integral position errors |
| $u_{pred} = g_p e_p + g_f e_f + g_i \int e_p$ | Current, future, and integral position errors |
| $u_{pred} = g_p e_p + g_f e_v + g_i \int e_p$ | Current, velocity, and integral position errors |
| $u_{pred} = g_p e_p + g_f e_v + g_f e_f$ | Current, velocity, and future position errors |

Error signals are denoted $e$ with subscripts indicating type (e.g., $e_p$ is current position error)
e.g., $e_p = x_{position,cursor} - x_{position,target=k}$

When there are multiple goals, and each target has different controllers gains, inferring a decision variable requires inferring how a weighted mixture of targets drives subjects’ control actions. To do this, we used ideas from control theory to develop a method for decomposing the subject’s instantaneous control (i.e., acceleration) into distinct contributions from each target. This previous work has shown control in a nonlinear cost landscape – when there are multiple targets and thus minima in the landscape – can be approximated by a compositional mixture of linear controllers, and the mixing weights in optimal settings are given by the relative cost/value of individual controllers (Todorov, 2009; Dvijotham and Todorov, 2012). Next, we state the general problem and connection to common mixture models, then our approach to estimating this quantity from behavioral data.

At every time point, we want to decompose the subjects measured control signal (𝑢_*obs*_(𝑡) = acceleration) into the proportion of their control signal attributable to individual prey (K) targets (𝑢_*k*_(𝑡)).

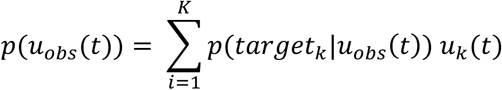

These controls are weighted by the posterior probability of them following a target, given their control signal. Estimation of these weights implicitly defines a multinomial posterior probability over K targets given the control signal. This posterior is the same form as the familiar gaussian mixture model (Murphy, 2022). Posterior latent target probabilities are then computed via Bayes rule:

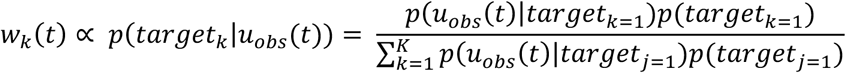

This shows that estimating 𝑤_*k*_(𝑡) will implicitly define the integration of an observed data likelihood and categorical prior. Because solving for the weights is a convex optimization problem, we will explicitly optimize (see Model parameterization) the controller weights 𝑤_*k*_(𝑡):

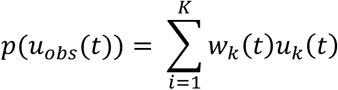

A final note worth mention is how optimizing 𝑤_*k*_(𝑡) (ie., target posterior) as a mixture model mixture approach is different from hidden Markov models. Mixture modeling applies soft-assignment or weighting and, as such, using 𝑤_*k*_(𝑡) to explain subject’s control signal allows blending of controllers and smooth transitions between them. Hidden Markov models use state decoding or hard-assignment, and effectively apply an argmax[𝑤_*k*_(𝑡)] to choose a singular controller. In the supplementary (section: model optimization) we show with simulations using a soft-assignment picks up both cases, where switching is either gradual or binary between targets; thus, using soft-assigned weights will correctly detect the target pursuit and switching strategy for either blending of controllers (soft-assignment) or discrete controller choices (hard-assignment).

#### Model parameterization

To infer an estimated 𝑤_*k*_(𝑡) time-series per trial, we estimated maximum a posteriori parameters from behavioral data, including the parameters of 𝑤_*k*_ ([𝜙, 𝜓]) and controller gains (𝑔_*k*_ = 𝜃). Both sets of parameters are explained below. Optimization of these parameters proceeded by minimizing the negative log posterior:

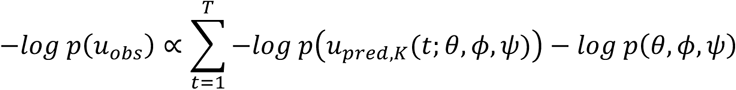

The first term on the r.h.s. is the negative log likelihood (priors are second). 𝑢_*pred*_ is the model predicted control signal that is determined by the mixture of controls across targets:

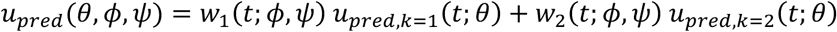

#### 𝑤_*k*_ parameterization

Mixture weights (𝑤_*k*_) were parameterized by implicitly modeling time dependent logits with (n=30) radial basis functions equally-spaced over each trial’s time. Basis parameters included weights (𝜙) and a shared width parameter (𝜓). We chose to implicitly parameterize 𝑤_*k*_ with basis functions as it allows a flexible and smooth parametrization of transitions between target control blending. Practically, 𝑤_*k*_ are defined though a softmax over the logits (eq. Xand X). Logits are computed by multiplying basis functions by their optimized parameters.

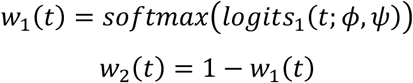

Note, this approach is readily extended to the case of K > 2 targets by constraining one target’s logits to 1 and increasing the basis parameters to n*K-1.

#### 𝒖_𝒑𝒓𝒆𝒅,𝒌_ parameterization

For generating predicted controls per target, 𝑢_pred,k_, we tested different controller types and estimate their gain parameters per controller (𝑔_*k*_). We model the predicted subject’s states (𝑥_*pred*_ ={positions, velocities}) and controls as a linear dynamic system mapping joystick input to the cursor. Joystick position controlled the cursor velocity, and therefore control signals from the subject (𝑢_*obs*_) are acceleration, which are composed of the controls from distinct targets (𝑢_*pred,k*_). These controls and states step the predictions forward and generate a new set of control predictions in each optimization iteration. The linear system describing the subject’s cursor is:

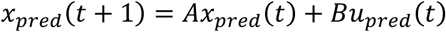

Where **A** and **B** are known system parameters (𝛥𝑡 = sampling rate (1/60 Hz)) that determine the mapping from joystick to subject cursor dynamics.

*A* = [10 Δ*t* 0; 0 1 0 Δ*t*; 0 0 1 0; 0 0 0 1], *B* = [0 0 Δ*t* 0; 0 0 0 Δ*t*]

Because the subject’s control strategy for a given target is unknown, we must also fit the mixing weights using different combinations of controller classes that could generate responses. We considered controllers that use different combinations of error signals to drive target pursuit. Model classes included either proportional or integral errors, where the error signals were of different classes described below, including feedback and feedforward predictive controller errors. All model classes are listed in Table 1.

Note that we also included models with future position errors. This controller class allows the subject’s predicted controls to be driven by feedforward signals based on the target’s predicted (1-step ahead) position. The error signal encodes the difference between the subject’s current (t) position and target’s future (t+1) position. Target future position was computed using the forward kinematics of the target:

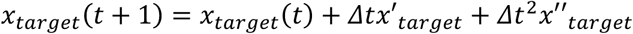

#### Model loss priors

The parameter priors for the loss are denoted as:

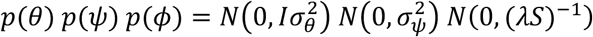

The prior over controller gains (𝜃) was a zero-mean normal distribution, with identity variance – constraining gains independently. A similar prior was used for the basis function width (𝜓). For both priors, increasing 𝜎^2^ decreases the regularization of the prior. For basis function weights (𝜙), we wanted to penalize deviations from smoothness. For this, we used a gaussian laplacian prior (multivariate gaussian) often used in spline models to penalize the wiggliness of functions (Wood, 2017). This prior uses the inverse precision matrix with a regularization coefficient (𝜆𝑆)^−1^. 𝑆 is a penalty matrix defined as the 2nd order difference between adjacent weights, penalizing the curvature:

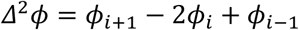

#### Model optimization

For parameter estimation, we used trust-const in scipy.optimize. During each parameter update, the current parameter set (𝜃, 𝜙, 𝜓) is used to simulate the predicted dynamics and compute the loss. Loss Jacobians and Hessians were computed automatically using Jax and parameters were iteratively optimized by maximizing the negative log posterior. To ensure positively constrained controller gains (𝜃) and basis width (𝜓) we optimized their log, and used a softplus transform. Each trial was optimized separately. For each optimization run, we fit a specific controller class (same for both targets). This was repeated 4 times for each controller class, with different parameter initializations, and the one with best loss was retained for cross model performance comparisons (next section). This was repeated for each of the seven model classes (Table 1).

#### Quantifying model performance and computing mixture

After optimizing parameters and retaining the best fit for each model class (and each trial), we computed the evidence lower bound (ELBO) for each model using a Laplace approximation to the parameter posteriors (see Murphy, 2022). The ELBO provides an estimate of lower bound on the marginal log-likelihood of each model, balancing model fit and penalizing the number of parameters.

The next step is computing a mixing trajectory (𝑒. 𝑔., 𝑤_1_) for each trial that marginalizes over each model, while accounting for each model’s uncertainty in predicted parameters. Because the model probability is quantified via the ELBO, we can use these ELBOs to compute posterior probabilities for each model.

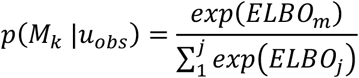

We use these model probabilities to perform Bayesian model averaging to obtain a singular 𝑤_1_ trajectory. In short, the 𝑤_1_ trajectory around the MAP parameters for each model class is weighted and summed. Computing 𝑤_1_ through model averaging accounts for model uncertainty by marginalizing over the generating models, and this is the target blending estimate we use for downstream analysis.

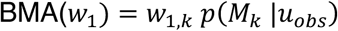

#### Comparison of different control mixing strategies

We also want to compare a smooth controller blending model to two other possible controller selection models that could drive behavior. We can use the same model fitting framework as above. For one comparison model, we fit a model that assumes subject’s choose a single controller at an given moment. This is equivalent to argmax model over individual target controllers, is nested within the smooth blending model, and is akin to how an HMM would choose the generative controller.

Comparison of this model to the smooth blending effectively tells us how much of behavior is explainable by a continuous mixing versus discrete selection strategy. This argmax model is formulated for the predicted control as:

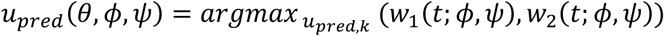

Another strategy that subjects could use is blending inputs with a single controller. Note that this control strategy is not nested within the smooth controller blending model, as control signals produced by multiple controllers versus one produced by weighting multiple inputs are not related through any linear transform (Supplementary note X). The input blending model is formulated using a single set of controller gains rather than distinct gains per target, and weighting the inputs:

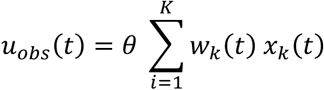

Both comparison models were fit using the same approach as the controller blending model. Bayesian model selection with the ELBO was used to select the best fitting model for each trial. For each subject, we computed the mean posterior probability of each model type for group level model selection.

### Behavioral analysis

#### Finding behavioral switch points

To separate instantaneous moments into pursuit versus change-of-mind points between targets, we first searched each trial to find points where 𝑤_1_ = 0.5. These points denote crossings between targets. (There is necessarily some element of estimation in the process of detecting switches, and, due to the inherent variability in human behavior, our algorithm necessarily has, at some point, an arbitrary cutoff.) We excluded any putative switch points that occurred within 325 ms of the trial start time, under the assumption that these likely represent a correction to an initial choice rather than true pursuit switching or change-of-mind. Using the remaining potential switch points, we created a window of -250 to +250 ms around these points. The switch trajectories were then categorized as falling into one of three categories. The main category (1) was those where a full switch was made between targets, the other two were either (2) partial switches or (3) were unclassifiable. To achieve this classification, we used a PCA across all switch trajectories from within a session. The input matrix for the PCA had switches index as rows and samples as columns (switches × samples). The 1^st^ principal component (PC) always captured the sigmoidal shaped full switch trajectory of interest. We computed the cosine similarity of this 1^st^ PC score and all switch trajectories and retained those with a similarity of > 0.9. We ran a PCA again on this subset, with the 1^st^ PC again producing the canonical sigmoidal trajectory. In this case, we retained those with cosine similarity of > 0.97. This threshold proved to be conservative enough to remove all partial switches, based on visual inspection. In total, an average of 35% of crossing-points were labeled as legitimate switches and forwarded for further analysis. Switch onset and offset timing were determined using a linear fitting change-point algorithm that looked for changes in mean and slope of the trajectory. To find the switch start, we searched backward in time from the crossing of 𝑤_1_ = 0.5. Switch ends were found by searching forward in time. Note that if two or more switches occur in the defined time window, their combined trajectory no longer resembles a clean sigmoidal transition. Because our PCA based classification relies on cosine similarity to the first principal component (which captures the prototypical sigmoid shape), such multi switches events would yield low similarity scores and would not be labelled as “full switches”. In our exploration of the data, we find that such double switches are very rare, so our approach is fundamentally conservative.

#### Switch statistics

The factors underlying the propensity to make target switches were modeled using a Bayesian binomial mixed effects model implemented in statsmodels in python (Seabold and Perktold, 2010). We used this approach to allow modeling all participants together, while accounting for individual variation in predictors. Predictor significance was determined using the 95% posterior credible interval. In terms of predictors, we asked if switch probability was sensitive to the time elapsed in trial, the difference in prey reward, and several prey distance variables. Specifically, we included the mean distance to the prey being pursued and the mean distance of the prey switched to, where the mean is calculated in the 100 ms leading up to a switch. We also included a factor to determine the mean rate of change of distance (distance closure) between both preys, too. The distance closure captures whether either prey getting closer or farther away (on average) altered switch probability. Each of these variables was centered by using their median values extracted from trials in which there were no switches. This allows interpreting the coefficients in terms of changes with respect to control, no-switch trials.

### Neural analysis

#### Single neuron encoding of task variables

To assess how neurons are tuned to task variables, we employed a linear-nonlinear Poisson model as a Generalized additive model (GAM; Balzani et al., 2020; Wood, 2017). Specifically, we aimed to quantify how neurons were tuned to subject speed, relative prey distance, relative prey speed, relative time in trial, prey reward difference, and the model-based blending variable. As our main goal was estimating whether neurons exhibited disentangled (linear) tuning to 𝑤_1_ or multiplexed with prey value, we included both 𝑤_1_ and its interactions with prey value as predictors. The GAM was fit with a penalized regression smoothing spline model (Wood, 2017). Penalized smoothing spline GAMs were chosen as they have been shown to offer good recovery of nonlinear neuron tuning, particularly in tasks where tuning is estimated with respect to continuously changing variables (e.g., speed; Balzani et al., 2020). Natural cubic regression splines served as a basis function. Cubic splines are convenient as a smooth interpolant basis because their only hyperparameter is the number of bases (𝐽; Wood, 2017). The spike counts 𝑦_1_ are fit in a GAM with univariate and interaction effects with basis functions 𝑏_*j,x_1_*_ (basis 𝑗 and variable 𝑥_1_). As an example, in a two-variable model the firing is modeled as:

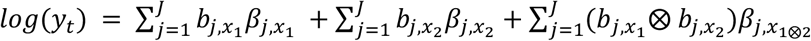

The first two terms are examples of single (marginal) variable tuning effects. The latter component encodes the interaction (nonlinear mixed selectivity) term as a tensor over the two univariate basis functions.

The model was fit in a Bayesian framework using stochastic variational inference in NumPyro. To estimate smooth and regularized tuning functions, we follow previous implementations (Balzani et al., 2020; Wood, 2017) and implement two types of regularization. Specifically, regularization was implemented through two types of priors on model coefficients that (1) control for basis wiggliness and enforce smoothness across basis functions, and (2) force sparse sets of bases similar to an L1 regularization. Combining these two effectively performs model selection by both smoothing across bases (1) and (2) forcing bases that don’t contribute to the model likelihood to be zero. All marginal and interaction terms were included in a single model and subject to the same smoothness and shrinkage priors. As a result, interaction terms that were not supported by the data were automatically shrunk toward zero, yielding an effective main-effects-only model when appropriate. These priors are implemented for basis coefficients as a multivariate normal distribution, for a given variable as:

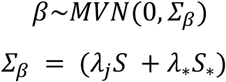

The smoothing/regularization terms (𝜆) are treated as random variables in the Bayesian approach, with Half-Cauchy priors. This allowed estimating separate smoothing terms per univariate and tensor model term directly from the data, without cross-validation. 𝑆 encodes a 2^nd^-order difference penalty between basis coefficients, inducing smoothing as a Laplacian graph prior that is used to construct the coefficients prior covariance. We point the reader to Wood (2017) for more specifics on how 𝑆 and 𝑆_∗_ are constructed.

To fit the models and approximate the posterior distribution of parameters, we employed a variational inference approach with a Gaussian guide. Variational parameters (mean and covariance) were optimized by minimizing the negative evidence lower bound (ELBO). This approach balances computational efficiency with parameter uncertainty estimation. All models were optimized using Adam (learning rate: 1e-3), each run for 10,000 steps.

To further determine coefficient significance, 99% posterior credible intervals were used to evaluate whether they included zero. Those variables with intervals non-overlapping with zero were retained. To determine whether neural activity was reliably modulated by task variables, we compared the full model against a baseline intercept-only model using the expected log predictive density (ELPD) estimated via WAIC (Gelman, 2013). The resulting WAIC model weight was used as a neuron-level model selection criterion, and only neurons for which the full model was favored over baseline were retained for subsequent analyses.

#### Fisher information of 𝒘_𝒕_ tuning

We computed a cross-validated and denoised estimate of fisher information using neuron tuning to 𝑤_*t*_. We computed tuning curves for each neuron in twelve bins. We used 5-fold cross-validation when computing the tuning curves. Tuning curves from the training set for each brain region were reduced using principal components analysis, retaining enough dimensions to account for 90% of variance. Fisher information was computed using PCA projections of the test set, and taking the Euclidean distance between adjacent levels of 𝑤_1_ of these projections (Kriegeskorte and Wei, 2021). To test for significant information, we created a null distribution for permutation testing by permuting 𝑤_*i*_ labels during calculation of the tuning curves. This distribution was created by repeating this process 1000 times, and the p-value was computed by comparing the mean actual 5-fold information to the permutation distribution.

#### Demixed principal components analysis

To examine the low-dimensional population dynamics related to behavioral controller blending, we performed dimensionality reduction on neural data using demixed principal components analysis (dPCA; Kobak et al., 2016). Using dPCA, we can extract linear subspaces of population activity that maximize the variance specifically related to separating levels of blending. In effect, this approach allows us to determine whether the population has linearly decodable representations of blending.

Because our aim was assessing whether neural population dynamics closely tracked the behavioral blending variable, we focused on time windows in which there was an explicitly consistent blending trajectory: change of mind between prey targets. To decompose the neural data, we first sorted all switch points, during a trial, for a given subject, and within a session, into those that switched from a low to high or high to low value target. Under the assumption that blending differs in a mirrored fashion (high to low versus low to high 𝑤_*t*_), across time, for these switch directions, we can take the decoding subspace as the dynamic subspace for 𝑤_*t*_.

We fit the dPCA model to firing rate tensors as pseudo-populations that included all subjects. The model included the effects for 𝑤_*t*_, prey reward difference, and their interaction. Each fit was done separately for ACC (n = 254), Hippocampus (n = 546) and OFC (n = 103). The fitting and significance testing procedure was carried out by performing 1000 repetitions randomly sampling (stratified monte-carlo) the data and holding-out 5% of trials (balanced between switch directions). A dPCA model was then trained using a fixed regularization of 1×10^-5^. Firing rate test-sets were then projected along the top decoder dimension for the switch direction. We took the decoded switch direction to be the minimum distance of the projected test data to the projected training data. This classification was computed for each of the 30 time-points in the switch window. Second, we established significance thresholds by repeating the above procedure, but randomly swapping trial labels to create a null accuracy distribution. We created the null distribution by repeating the 1000 iterations 50 times. For each of the 50 sets, we computed the null decoding accuracy. We determined that decoding was statistically significant when the decoding accuracy was greater than the null for 100% of the 50 sets.

To visualize how neural population activity varied across blending states and reward conditions, we performed an additional dPCA analysis in which neural activity was binned by the behavioral blending variable 𝑤_*t*_. This analysis was performed separately for trials with equal prey value and trials with different prey value. Neural firing rates were aligned to switch events. For each condition, firing rates were grouped into bins according to the instantaneous value of 𝑤_*t*_ and averaged to form a population activity matrix spanning time and blending state. We fit the dPCA model to firing rate tensors as pseudo-populations that included all subjects. We applied dPCA with marginalizations over blending state and its interaction with time, and projected population activity onto the leading components associated with these terms. The resulting low-dimensional activity patterns were visualized as heatmaps illustrating how neural dynamics evolved across time and blending state, separately for hippocampus, ACC, and OFC.

#### Clustering ramp

We clustered firing during the target switches to assess whether classes of single neurons also exhibit ramping behavior. To obtain a meaningful clustering based on latent neural dynamics and potential, we first fit a linear dynamic system to trial-averaged, gaussian smoothed (FWHM 60 ms) firing rates using the ssm package in Python. In addition, the models were fit to both switch directions (high to low and low to high value prey) simultaneously, giving them shared parameters but different inputs. We used the normalized relative target distance as inputs to the model. This approach to clustering accounts for similar latent dynamics evolving under different inputs. The number of the latent state dynamics for each model were optimized by computing the loss (ELBO) from a held out test in 5-fold cross-validation. This was done separately for each brain region. We found three latent states were suitable for describing each area.

Clustering was then performed on the firing rate embedding matrix C using K-means. The C matrix links latent states to firing rates. The number of clusters was optimized using the silhouette score, separate for each brain region. We chose clusters to retain based on those that exhibited significant differences in activity between time-locked segments with switched and control segments without. Significance was determined by permuting (1,000 times) the mean clustered firing rate between the actual switches and controls.

## Supplementary Material

**Figure S1.**
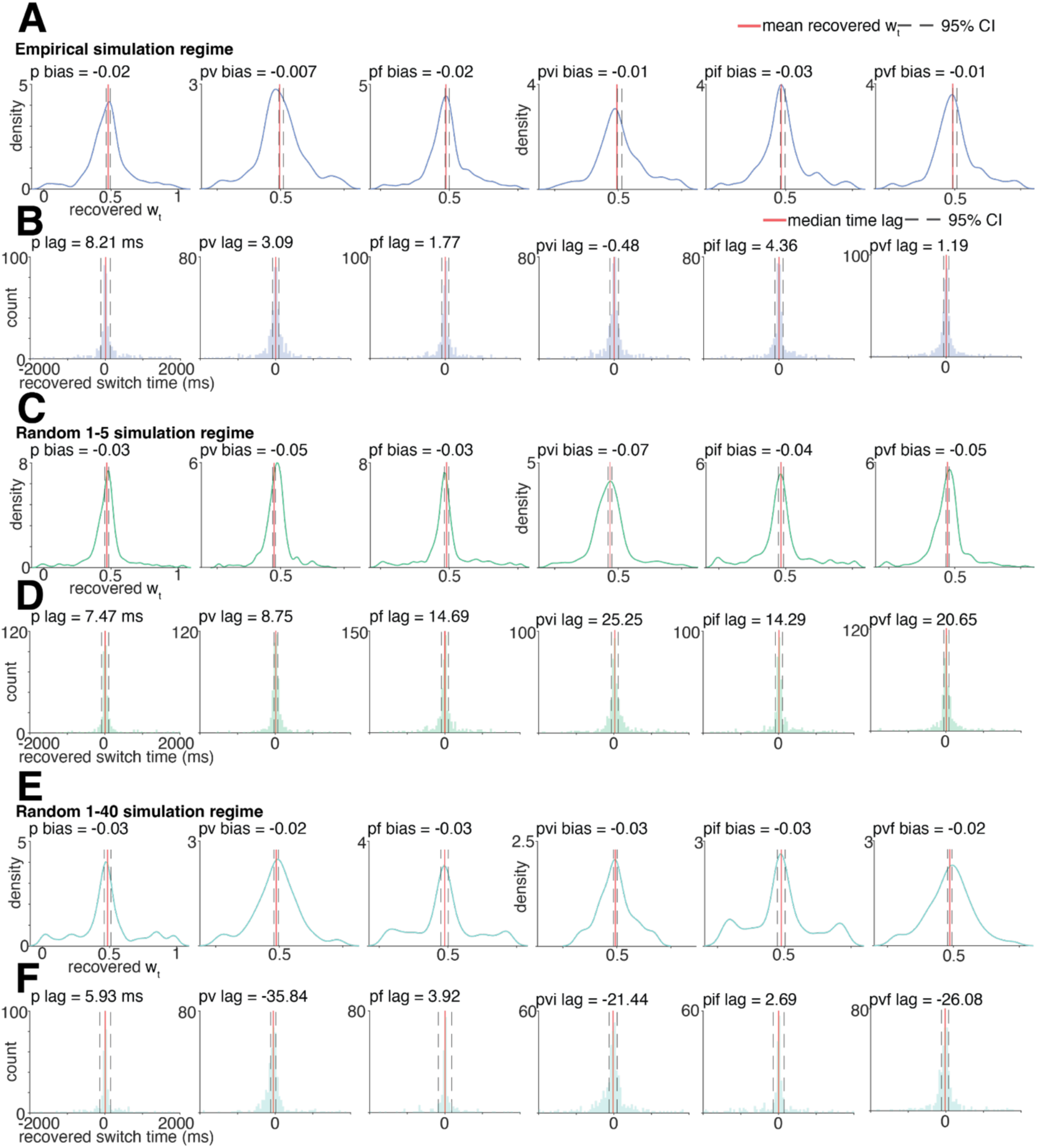
Model recovery of w_t_ value and time at switch point (w_t_ = 0.5, t = 0 ms). **A**) Empirical simulation regime - distribution of recovered w_t_ values at true switch value = 0.5, quantifying bias in switch magnitude recovery across controller models. The red line represents the median recovered value, dashed lines the 95% confidence intervals. (**B**) Distribution of temporal lag between recovered and true switch times for w_t_ = 0.5, quantifying timing accuracy of switch detection across controller models. The red line represents the median recovered value, dashed lines the 95% confidence intervals. (**C**) - (**E**). As in panel A, but for the empirical simulation regimes. (**D**) - (**F**) As in panel B, but for the empirical simulation regime.

**Figure S2.**
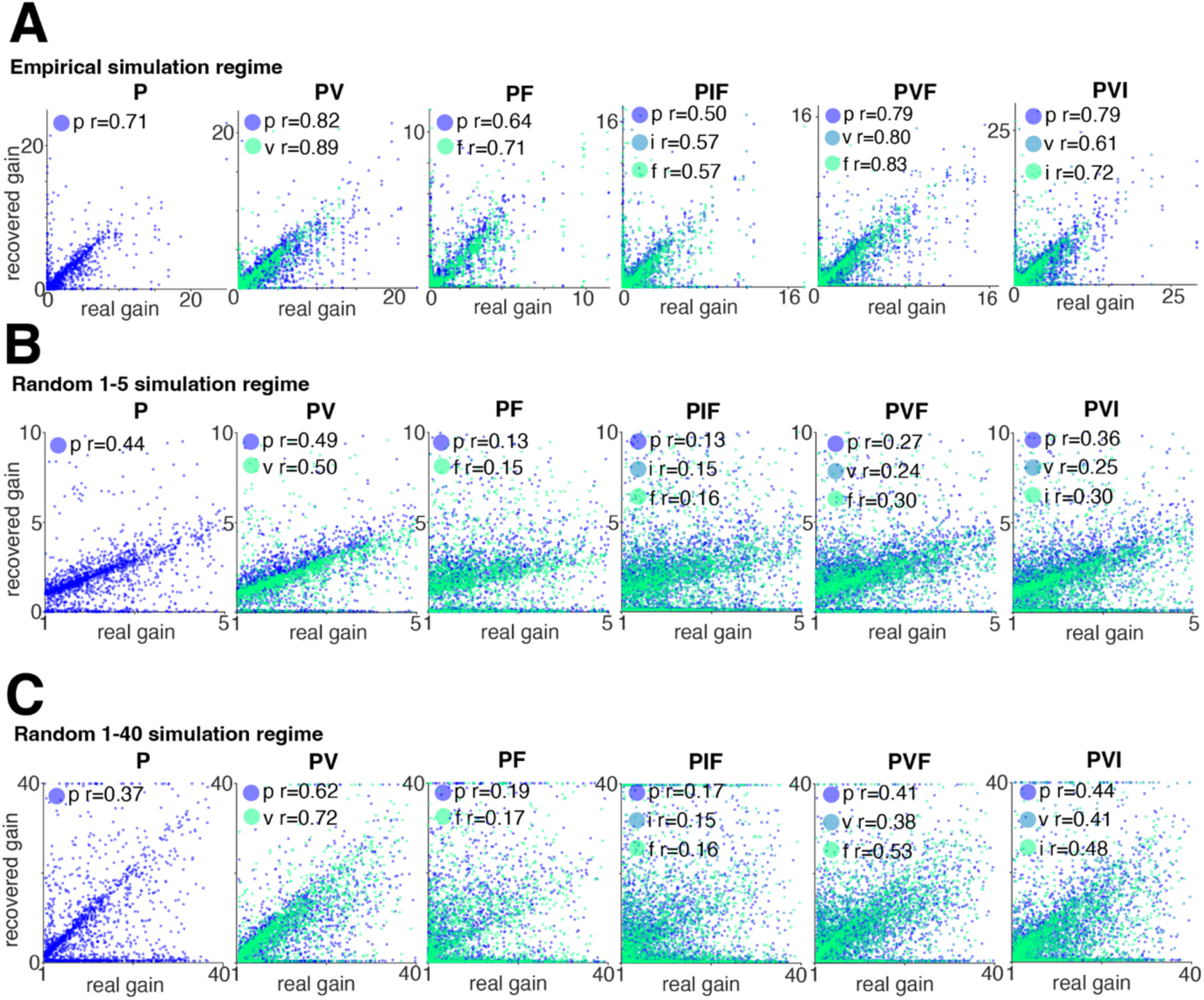
Model recovery of controller gains across controller classes. (**A**) Scatter plot illustrating the correlation between the true gains and the gains recovered from the model. Each column represents a controller class. (**B**) - (**C**). As in panel A, but for the simulation in the random regime.

**Figure S3.**
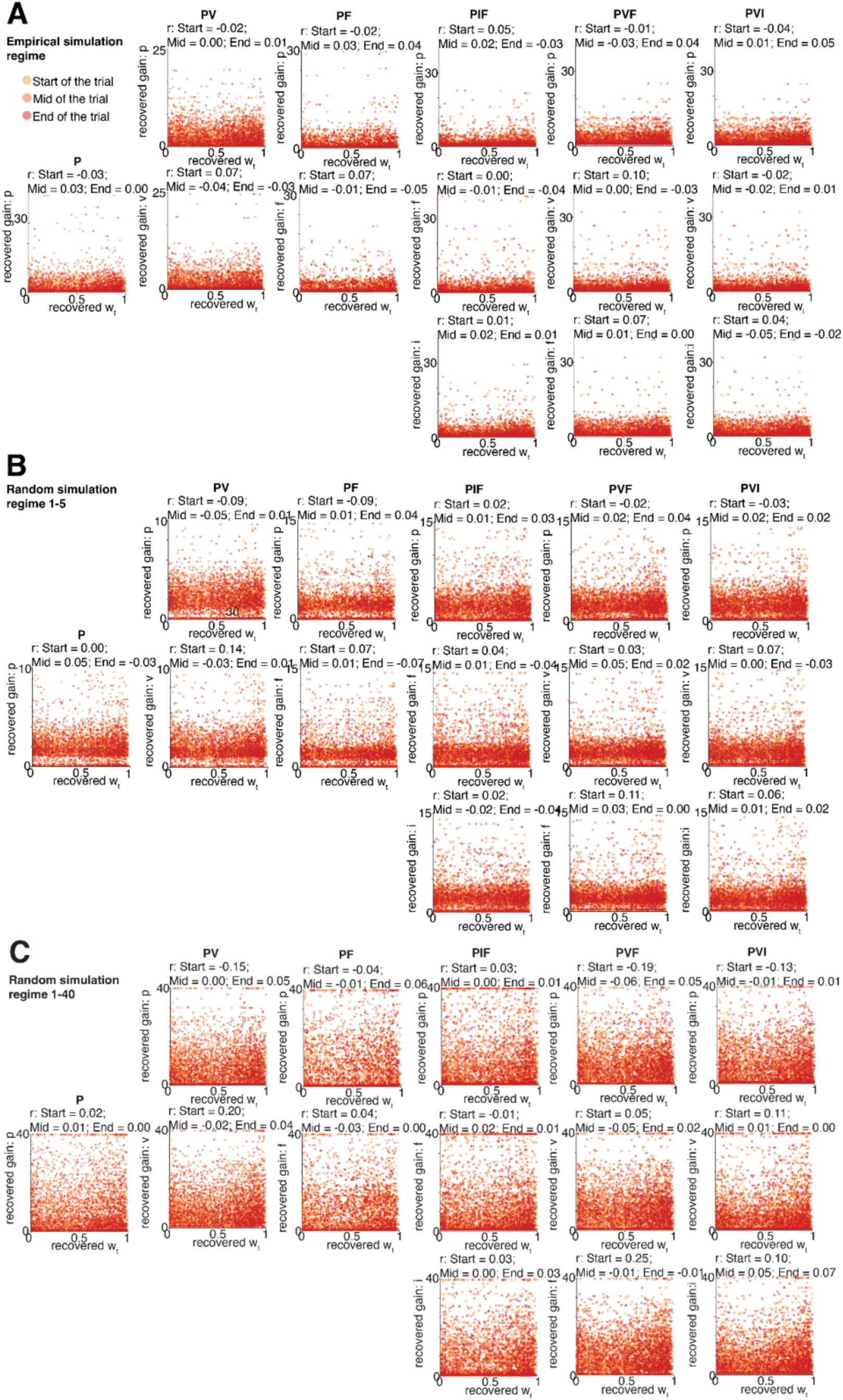
Model recovered controller gains do not correlate with w_t_ recovered trajectories. (**A**) Scatter plot illustrating the lack of correlation between the recovered controller gains and the recovered w_t_ trajectories, each column represents a controller class (empirical simulation regime). (**B**) - (**C**). As in panel A, but for random simulations.

**Figure S4.**
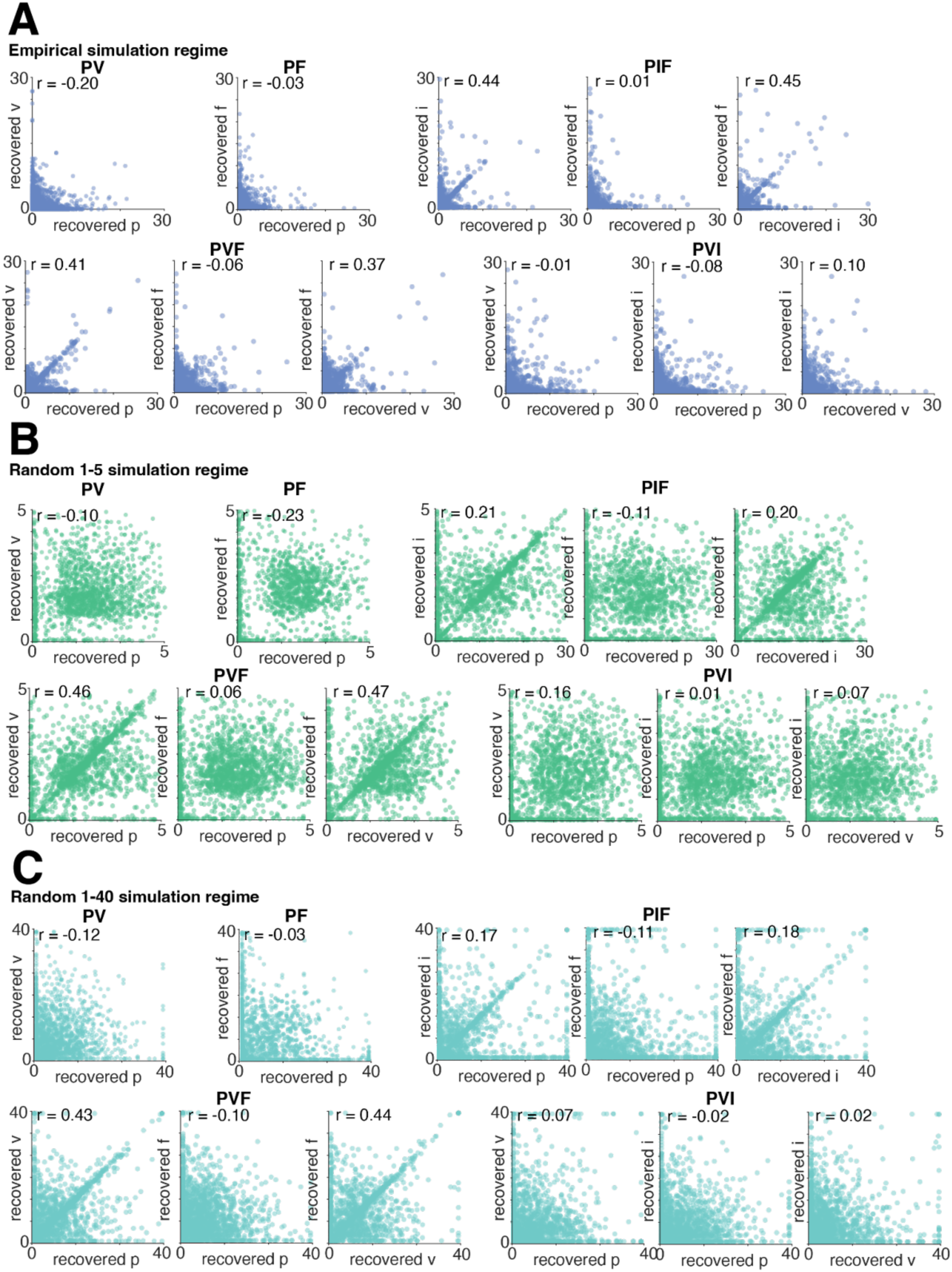
Model recovered controller gains largely do not correlate with each other. Scatter plot illustrating the lack of correlation across most of the recovered controller gains across simulations.

### S5. Mixture of controllers model fitting accuracy and ground-truth recovery

To test the validity of our modeling approach using basis functions and controllers, we simulated ground-truth data with a known w_t_ that we designed. To cover the space from simple to complex w_t_ profiles, we considered five different types of w_t_. To simulate data, we sampled 20 random trials across both subjects’ sessions, with 5 of each having an approximate length of 0.91 sec, 1.45 sec, 2.83 sec, 4.1 sec long. In the main text, we fit across a range of number of bases and model-average. We chose a fixed number of bases in simulation to fix the complexity of the model, as increasing basis numbers will always increase fit quality.

As a first test, we compared whether different controller types (see Table 1) could be recovered as the best fitting model compared to a baseline of just a position controller (**Figure S5A**). The posterior model probabilities indicated that for all controller classes we could recover the true generative controller model on average better compared to a baseline (all probabilities > 0.5 for the generative model). We then performed a full confusion analysis across all three simulation regimes. For each generative model defined by the controller class, we simulated datasets using known parameters, fit all the candidate models to each dataset, and selected the best fitting model using the ELBO. We obtained a full confusion matrix quantifying how reliably each generative controller class could be distinguished from the others (**Figure S6**). We found that recoverability was statistically significant across simulations for controllers p, pv and pvi (binomial test, p < 0.0001 in all cases), but not for the remaining classes. Importantly, the pv model, which turned out to be the best-firring model across participants, was consistently and reliably recovered (43% correct; 95% CI [0.40 0.47], p < 0.0001), confirming that this class can be identified despite partial overlap between other models.

We next examined how well we could recover the generative kinematics for each controller model class. In **Figure S5B** we show the average correlation between predicted and actual positions for 20 simulated trials per controller model type. We get good recovery across all model types, with an average correlation of 0.83 across model classes. The quality of inferring ground-truth w_t_ was similar to positions. The correlations between the true and predicted w_t_ are shown in **Figure S5C**. Across the generative models, we get an average correlation of 0.87. These correlations are averaged over different levels of w_t_ complexity, which was controlled using a scaling parameter for a generating gaussian process model. In **Figure S5D-I**, we show example fits of the simulated and recovered w_t_ for each class of generative models. Overall, these simulations and analysis indicate that our method did a good job at detecting the correct model class (**Figure S5A**), as well as recovering the actual latent target specific controls (**Figure S5B**) and w_t_ (**Figure S5C**).

**Figure S5.**
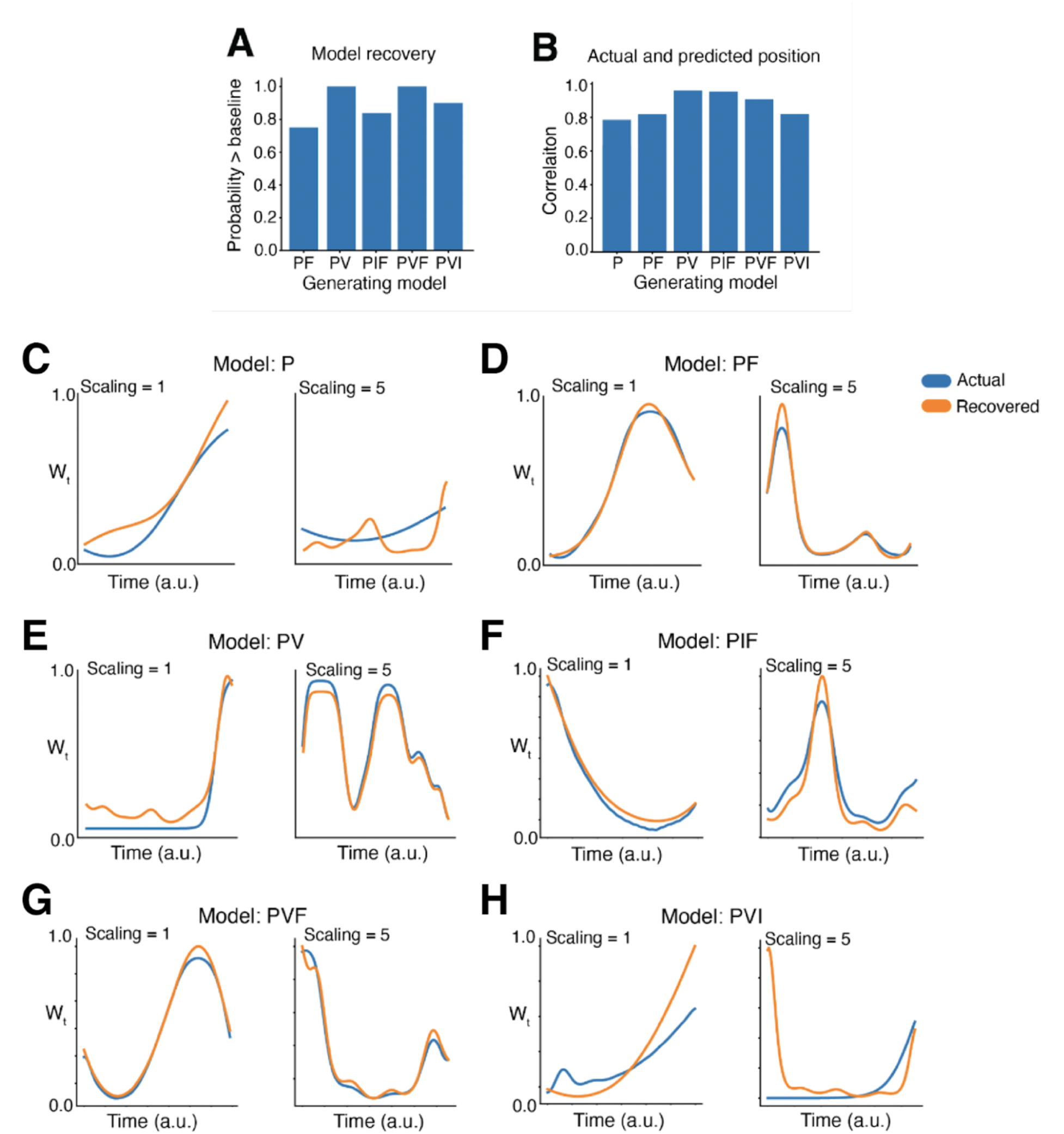
Model recovery of w_t_ and controllers. (**A**) Probability of choosing the generative model as being better than a baseline, position only controller model. Higher values indicate better recovery. (**B**). Correlation between true and predicted controller position time series based on fit of generative model class. Higher correlations indicate better recovery, and each bar is the average of 30 runs. (**C**) - (**H**) examples from simulations and fits showing the generative and the predicted model recovered w_t_. Each set of plots is a different generative model. The left columns in each (scaling = 1) are those with simple generating w_t_, and the right columns are more complex w_t_ (scaling = 5). The scaling is the scale parameters used in a gaussian process generator.

**Figure S6.**
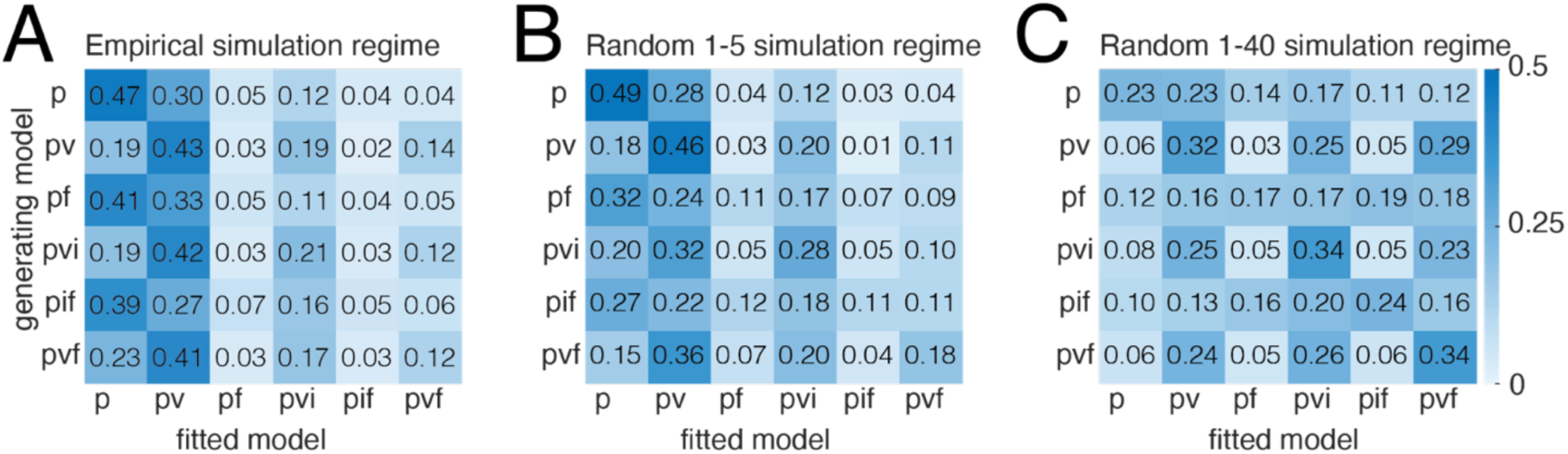
Full model recovery confusion matrix. **(A)** Confusion matrix quantifying how reliably each generative controller class could be distinguished from the others

### S7. Proof of controller model non-equivalencies

We outline a simple algebraic proof here that two of the considered model classes – mixing of controllers versus mixing of inputs – are not related by any linear transform, making them constitute different behavioral model classes. We formulate this argument for a single time-point, input feature and controller dimension. An assumption we make is that controller gains are at steady state, as formulated in an infinite horizon or average cost scenario, ie., controller gains are not state or time-dependent. This problem is formulated is as follows:

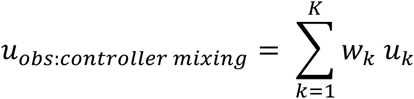

Which represents the controller blending model, where control signals for two controllers are specified with gains (𝐿_*k*_):

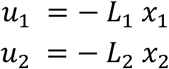

Using this, it is convenient to rewrite this as:

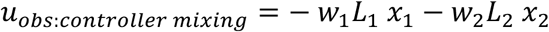

For comparison, we create the same setup for input mixing models:

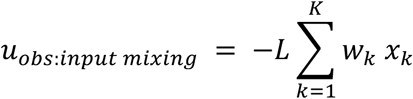

Rewriting in expanded form:

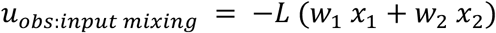

Now testing for equivalence across models, we find that are only equivalent when the controller gains are identical:

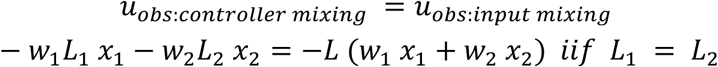

Thus, the two models can only be seen as equal when the gains of both controllers are equal.

**Figure S8.**
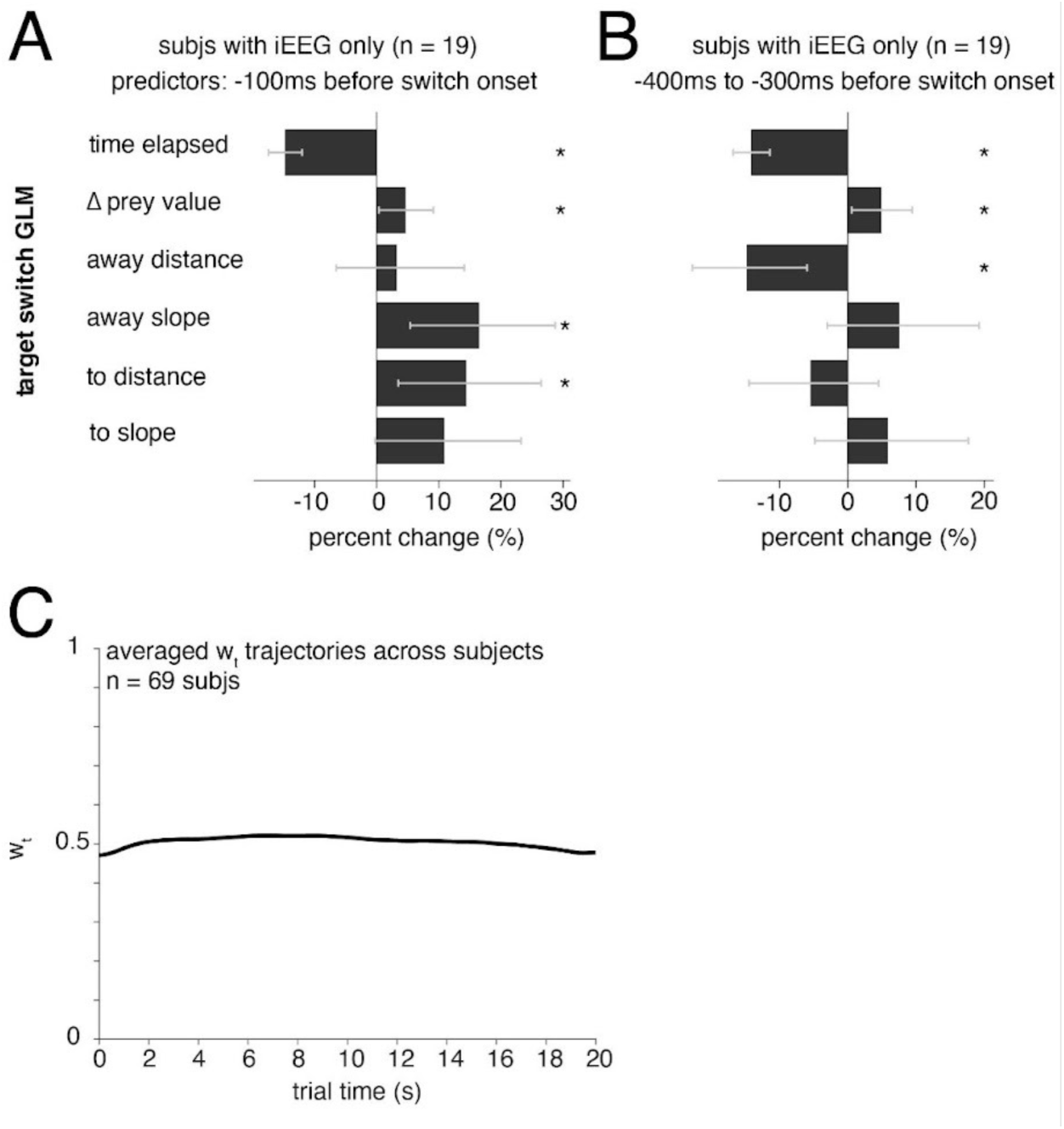
Behavioral predictors of target switching and average controller dynamics. **(A)** Target switch GLM including subjects with iEEG only. Predictors computed using the mean variable (i.e., time elapsed) 100ms before the switch onset. (**B**) Target switch GLM including subjects with iEEG only. Predictors computed using the mean variable (i.e., time elapsed) between -400ms and -300ms from switch onset. (**C**). Averaged w_t_ trajectories across all the participants.

